# Auto-methylation of the histone methyltransferase SetDB1 at its histone-mimic motifs ensures the spreading and maintenance of heterochromatin

**DOI:** 10.1101/2025.01.21.634156

**Authors:** Qing Tang, An Zhang, Michael Sullivan, Katalin Fejes Toth, Alexei A. Aravin

## Abstract

Heterochromatin plays a critical role in nuclear organization and the regulation of gene expression by directing 3D genome organization, regulating lineage-specific gene expression, and ensuring the repression of transposable elements and endogenous retroviruses. Functionally and structurally distinct chromatin domains are defined by the so-called histone code, which consists of combinations of post-translational histone modifications deposited by “code writers” and recognized by “code readers.” The primary mark of heterochromatin, trimethylation of histone H3 at lysine 9 (H3K9me3), is deposited by histone methyltransferases, such as SetDB1, and serves as a binding platform for readers, most notably HP1 family proteins.

Using a reporter system to monitor the dynamics of heterochromatin establishment and maintenance, we demonstrated that transient tethering of HP1 triggers the SetDB1-dependent establishment of stable heterochromatin. This finding indicates the presence of a feedback mechanism wherein the reader of the H3K9me3 mark recruits the writer. We further discovered that the genetic interaction between SetDB1 and HP1 is mirrored by a direct physical interaction. This interaction requires the auto-methylation of two conserved histone mimic motifs located in unstructured regions of SetDB1. HP1 binds these SetDB1 motifs using the same molecular interface it employs to recognize the modified histone tail.

Our findings show that SetDB1 auto-methylation is essential for the spreading and stable maintenance of heterochromatin. This includes its roles in processes such as X-chromosome inactivation and the negative feedback regulation of a large gene family encoding KRAB-ZNF transcriptional repressors. Thus, the primary heterochromatin mark is not limited to nucleosomes but is also deployed on the mark’s writer itself. This fosters a direct physical interaction between the writer and the reader, ensuring key features of heterochromatin: its spreading to establish extended domains and its stable maintenance through cell divisions.

## Introduction

Heterochromatin is a highly compacted form of chromatin typically found in regions of the genome that are transcriptionally inactive. Proper heterochromatin formation is essential for cellular function, as the loss of heterochromatin can lead to altered gene expression, genome rearrangements, and genomic instability (Allshire and Madhani 2018; Becker et al. 2016; Janssen et al. 2018; Peng and Karpen 2009). Based on its characteristics and functions, heterochromatin is classified into two types: constitutive heterochromatin, which is permanently condensed in all cell types, and facultative heterochromatin, which is dynamic and controls lineage-specific gene repression during development in a tissue-specific manner (Trojer and Reinberg 2007; Allshire and Madhani 2018; Oberdoerffer and Sinclair 2007). Facultative heterochromatin often contains genes that can be turned on or off, enabling flexible gene regulation in response to environmental cues or developmental signals (Oberdoerffer and Sinclair 2007). In contrast, constitutive heterochromatin is primarily associated with repetitive DNA sequences, such as simple repeats found around centromeres and telomeres, as well as transposable elements (TEs). It serves as a structural component of chromosomes, aiding in nuclear organization and maintaining chromosome integrity (Allshire and Madhani 2018; Saksouk et al. 2015; Bizhanova and Kaufman 2021). Interestingly, recent studies have revealed that features traditionally attributed to constitutive heterochromatin can also contribute to cell-type-specific gene expression profiles, blurring the distinction between the two heterochromatin states (Nicetto et al. 2019; Nicetto and Zaret 2019; Padeken et al. 2022).

Different chromatin regions are marked by chromatin modifications that, often in combination, specify the functional state of the underlying DNA. These modifications are deposited by “code writers,” enzymes that post-translationally modify histones or methylate DNA. Constitutive heterochromatin is characterized by di- and trimethylation of lysine 9 on histone H3 (H3K9me2/3), a modification deposited by H3K9-specific histone methyltransferases (HMTs). In mammals, six H3K9 HMTs have been characterized, each with distinct but partially overlapping targets, serving different purposes (Padeken et al. 2022). Of these, only SetDB1/2 and Suv39h1/2 can deposit the trimethyl mark, while G9a and GLP deposit mono- and dimethyl marks. Together, these enzymes silence pericentromeric and telomeric repeats as well as TEs (Padeken et al. 2022). Additionally, they silence many genes, with SetDB1 notably involved in X-inactivation (Keniry et al. 2016; Sun and Chadwick 2018; Minkovsky et al. 2014) and the repression of a large class of KRAB zinc-finger (KRAB-ZNF) transcription repressors, which themselves help target SetDB1 to TEs (O’Geen et al. 2007; Frietze et al. 2010; Tchasovnikarova et al. 2015; Wolf et al. 2015). Most chromatin marks function as binding platforms for “code readers,” regulatory proteins that either exert their regulatory function directly or recruit additional factors (Hyun et al. 2017). The hallmark reader of heterochromatin is heterochromatin protein 1 (HP1), a family of proteins that recognize H3K9me2/3 through their N-terminal chromodomain (CD) (Bannister et al. 2001; Lachner et al. 2001; Jacobs and Khorasanizadeh 2002; Nielsen et al. 2002) and mediate transcriptional repression via their C-terminal chromoshadow domain (CSD) (Hathaway et al. 2012; Yan et al. 2018). The CSD is responsible for HP1 dimerization, which not only generates an interaction platform for other heterochromatin modifiers (Lechner et al. 2000; Thiru et al. 2004; Hediger and Gasser 2006), but also plays an important role in chromatin compaction (Hiragami-Hamada et al. 2016; Azzaz et al. 2014).

Heterochromatin formation typically involves three phases: initial establishment, spreading along the chromosome, and epigenetic maintenance through cell divisions. Initial establishment depends on the specific recruitment of HMTs to genomic target loci. Since HMTs lack sequence specificity, their recruitment is facilitated by various mechanisms. HMTs are often guided to chromatin by non-coding RNAs, such as Ago-bound siRNAs in worms, plants and fission yeast (Ahringer and Gasser 2018; Henderson and Jacobsen 2007; Holoch and Moazed 2015); piRNAs in diverse animals (Czech et al. 2018); RNAs transcribed from major satellite repeats and telomeric repeats in mammals (Shirai et al. 2017; Johnson et al. 2017; Velazquez Camacho et al. 2017; Porro et al. 2014); or long non-coding RNAs such as Xist (Strehle and Guttman 2020). In addition to non-coding RNA guides, many targets are recognized by specific DNA-binding proteins that recruit HMTs. For instance, in mammals, SetDB1 is recruited to endogenous retroviruses through KRAB-ZNF transcription repressors, which have evolved to recognize specific retroelements (Schultz 2002; Wolf et al. 2015). SetDB1 has also been suggested to target a diverse array of loci via the HUSH complex, which is proposed to recognize long intronless RNAs, such as LINE1 retroelements, pseudogenes, and plasmid derived RNAs (Seczynska et al. 2022). Such mechanisms enable transcription-dependent but sequence-independent establishment of heterochromatin domains.

Many targeting mechanisms are restricted to specific genomic regions or sequences, from which heterochromatin spreads into neighboring regions to form large and stable transcriptionally inactive domains. These domains are often stably maintained over extended periods, multiple cell divisions, or even generations. Spreading and maintenance frequently occur independently of the original recruiting signal and rely on stabilizing interactions between HMTs and chromatin (Hall et al. 2002; Ragunathan et al. 2015; Tatarakis et al. 2023). Recruitment mechanisms in which the depositing enzyme is recruited by its own histone mark, termed the read-write mechanism, can create a self-propagating system where newly added marks attract more silencing factors (Ragunathan et al. 2015; Grewal 2023). This system reinforces the heterochromatic state and facilitates its spreading along the chromosome. In fission yeast, the HMT Clr4/Suv39h binds H3K9me3, enabling a read-write mechanism for heterochromatin spreading and maintenance (Zhang et al. 2008; Ragunathan et al. 2015). Similar mechanisms have been observed in mammals, where G9a and GLP bind mono- and dimethylated H3K9 through their ankyrin repeats (Collins et al. 2008), while Suv39h1/2 interact with H3K9me2/3 via their N-terminal chromodomains (O’Carroll et al. 2000; Melcher et al. 2000; Wang et al. 2012; Weirich et al. 2021). These interactions were also reported to enhance HMT catalytic activities and promote propagation of the modification state (Liu et al. 2015; Müller et al. 2016). In contrast to the other mammalian HMTs, the interaction of SetDB1 with H3K9me3 is indirect, mediated by its cofactor ATF7IP binding MPP8, a core component of the HUSH complex, which binds H3K9me3 via its chromodomain (Schultz 2002; Wang et al. 2003; Chang et al. 2011; Tchasovnikarova et al. 2015; Tsusaka et al. 2018). To date, it is not known whether these reader-writer interactions have any function in mammalian heterochromatin spreading and maintenance.

Over the past three decades, the chromatin field has focused on understanding the functions, readers, and writers of histone modifications. However, post-translational modifications are not exclusive to histones. Emerging evidence suggests that non-histone proteins can “mimic” histone motifs to mediate protein-protein interactions (Sampath et al. 2007; Chin et al. 2007; Tsusaka et al. 2018; Ferry et al. 2017; Marazzi et al. 2012; Tarakhovsky and Prinjha 2018; Chen et al. 2021). Influenza virus exploits histone mimicry to suppress host antiviral response by interacting with host histone “reader” using histone mimic motif (HMM) on the viral protein (Marazzi et al. 2012). HMMs have been identified in several non-histone host proteins as well, the first case was discovered in the HMT G9a (Sampath et al. 2007; Chin et al. 2007). These HMMs can be methylated by G9a itself and interact with the chromodomain of HP1 in a methylation-dependent manner (Sampath et al. 2007; Chin et al. 2007). Subsequent studies identified HMMs in several non-histone substrates of G9a through biochemical screens (Rathert et al. 2008; Tsusaka et al. 2018). For instance, the SetDB1 cofactor ATF7IP contains an HMM that mediates its interaction with MPP8 (Tsusaka et al. 2018). However, the functional roles of these methylated HMMs remain unexplored.

In this study, we uncovered a genetic and direct physical interaction between human SetDB1 and HP1. We found that HP1 binds SetDB1 using the same chromodomain interface it employs to bind the H3K9me3 mark. We identified two motifs in the unstructured regions of SetDB1 that mimic the histone H3 tail and are autocatalytically modified by SetDB1 to create a complete mimic of the H3K9me3 mark. Loss of methylation on these HMMs impairs SetDB1’s interaction with HP1. Using a reversible tethering reporter system, we demonstrated that the methylation of these HMMs is crucial for the spreading and maintenance of ectopic heterochromatin but does not affect its initial establishment. Furthermore, mutation of the SetDB1 histone mimic disrupted heterochromatin spreading over native SetDB1 genomic targets, such as KRAB-ZNF clusters and the X chromosome. Interestingly, histone mimic mutations (HMm) also resulted in the formation of ectopic heterochromatin in euchromatic regions, indicating that HMM-mediated SetDB1-HP1 interaction is essential for constraining H3K9me3 modification to canonical heterochromatin domains. Thus, our study identifies a direct interaction between HP1 and SetDB1 mediated by histone mimicry and demonstrates that this interaction facilitates the formation and maintenance of large transcriptionally silent heterochromatin domains across multiple cell divisions.

## Results

### Transcriptional repression and formation of stable heterochromatin induced by HP1 recruitment requires the histone methyltransferase SetDB1

To investigate the establishment and maintenance of heterochromatin domains, we developed a reporter system for temporally controlled recruitment of chromatin writers and readers. The EGFP reporter gene was integrated into the genome of HEK293 cells at a euchromatic locus marked by active histone H3K4me3 and H3K36me3, but lacking H3K9me3, the hallmark of heterochromatin (Fig. 1A). Upstream of the elongation factor 1α promoter (pEF) that drives EGFP expression, we incorporated five Tet Operator (TetO) binding sites for the reverse Tet repressor (rTetR), which binds only in the presence of doxycycline (Dox) (Urlinger et al. 2000). Stable cell lines were selected using a hygromycin resistance gene driven by the SV40 early promoter (pSV40), located more than 5 kb upstream of the TetO sites. The Dox-inducible system ensures precise temporal control for the recruitment and release of rTetR-fused proteins to the reporter. This setup allows monitoring of ectopic heterochromatin establishment, maintenance, and spreading (Fig. 1B). EGFP fluorescence, measured via flow cytometry, provides a quantitative readout of reporter expression in individual cells.

**Fig. 1.**
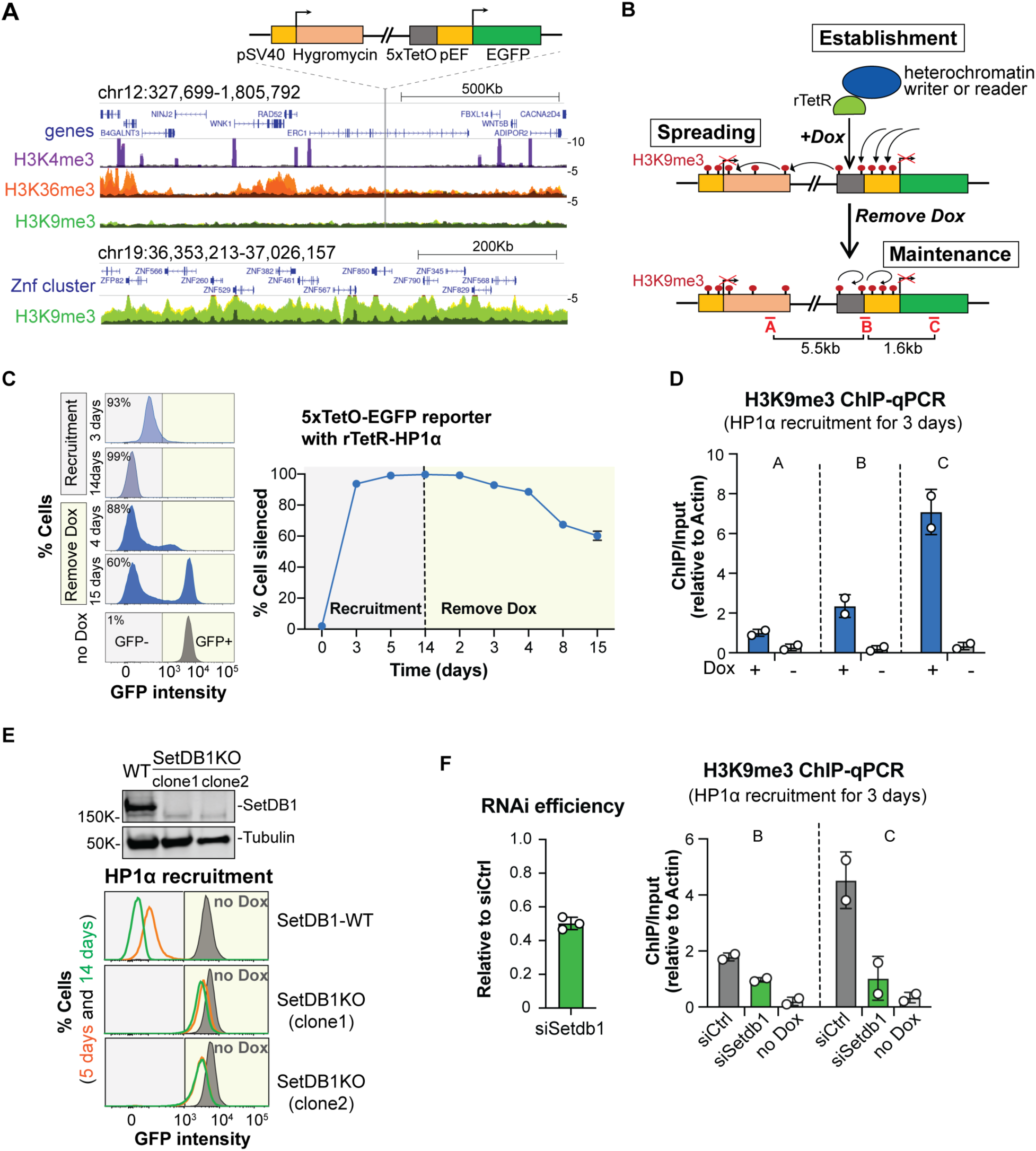
SetDB1 is required for heterochromatin establishment induced by HP1 recruitment. (A) Reporter architecture and characterization of its integration site. A reporter containing EGFP driven by the elongation factor 1α promoter (pEF), preceded by five Tet Operator sites (5xTetO), and a hygromycin resistance gene driven by the SV40 early promoter > 5kb upstream was site-specifically integrated into the genome of HEK293 cells. UCSC browser snapshot of ENCODE H3K4me3, H3K36me3, and H3K9me3 ChIP-seq tracks shows the euchromatic integration site. Two replicates of ChIP (colored) and input (gray) signals are overlaid. The H3K9me3 track on a KRAB-Znf gene cluster is shown as a reference for an H3K9me3-enriched region. (B) Schematic diagram of the inducible reporter system for studying heterochromatin. The rTetR-fused heterochromatin protein is stably co-expressed with the reporter and binds to TetO sites upon addition of doxycycline (Dox), and is released upon Dox removal. Analysis of transcriptional silencing efficiency and deposition and spreading of heterochromatin marks upon Dox supplementation and its removal after silencing establishment enables temporal evaluation of heterochromatin establishment, spreading, and maintenance. Red bars A–C denote amplicons used for qPCR analysis. (C) Artificial recruitment of HP1α leads to reporter silencing. GFP signal was measured by flow cytometry at different time points after rTetR-HP1α recruitment to the reporter. Dox was removed after 14 days of recruitment, and GFP signal measurement continued for an additional 15 days. The black line indicates the gating threshold to distinguish between GFP-positive (GFP+, yellow shading) and GFP-negative (GFP-, gray shading) cell populations. Dox-treated (blue) and control (gray) samples are shown. Right: Quantification of three biological replicates at indicated times. SD is plotted, but is too small to show for most points. (D) HP1α tethering induces H3K9me3 deposition over EGFP and the hygromycin resistance gene. H3K9me3 ChIP-qPCR was performed 3 days after HP1α recruitment. H3K9me3 ChIP-qPCR analysis was performed with and without Dox at three regions over the reporter as indicated in panel B. Dots correspond to independent biological replicates; bars indicate mean and SD. (E) SetDB1 is required for reporter silencing upon HP1α tethering. SetDB1 was knocked out in HEK293 cells that stably express both the reporter and rTetR-HP1α. Two single clones were isolated and validated by PCR (data not shown) and western blot (top). HP1α recruitment to the reporter was performed in SetDB1-WT cells and the two SetDB1-KO clones. GFP signals were measured by flow cytometry at day 5 (orange) and day 14 (green) after Dox treatment, or without Dox (gray). (F) SetDB1 knockdown decreases H3K9me3 deposition upon HP1α tethering. Control siRNA and siRNA against SetDB1 were transfected into cells stably expressing both the reporter and rTetR-HP1α. RNAi efficiency was determined 5 days post-siRNA transfection (left). HP1α was recruited to the reporter two days after siRNA transfection, and H3K9me3 ChIP-qPCR was performed 3 days later. Two regions are detected as described above. Dots correspond to independent biological replicates; bars indicate mean and SD.

To test whether the recruitment of the histone methyltransferase SetDB1, one of the primary writers of the H3K9me3 mark, induces heterochromatin formation, we integrated a transgene encoding SetDB1 fused to the rTetR domain into the reporter cell line. Recruitment of SetDB1 by Dox supplementation resulted in a progressive loss of EGFP reporter expression (Fig. S1A). As expected, repression was accompanied by the accumulation of H3K9me3 and HP1α (hereafter referred to as HP1) on the reporter, indicating that transcriptional silencing is linked to heterochromatin formation (Fig. S1B and C). Furthermore, the *de novo* deposition of H3K9me3 and HP1 following SetDB1 tethering extended >5 kb to the distal hygromycin resistance gene (Fig. S1B and C). Thus, recruitment of SetDB1 to the reporter initiates the formation of a heterochromatin domain, characterized by H3K9me3 and HP1 accumulation at and around the initiation site, and efficient transcriptional repression, which persists for several days after Dox removal from the media (Fig. S1).

Next, we recruited HP1 to the reporter, which also resulted in strong reporter repression (Fig. 1C). ChIP-qPCR revealed H3K9me3 accumulation on the reporter (Fig. 1D), indicating that HP1 tethering induces bona fide heterochromatin similar to that observed upon SetDB1 recruitment. Notably, the reporter remained repressed even after HP1 recruitment was terminated by Dox removal (Fig. 1C). Approximately 60% of dividing cells maintained complete reporter repression two weeks after HP1 recruitment ceased, demonstrating that established heterochromatin is stable through cell divisions, even without ongoing HP1 tethering (Fig. 1C). These findings suggest that HP1 is not merely a reader of the H3K9me3 mark but also provides a feedback loop that enables heterochromatin formation, likely by recruiting histone methyltransferases. This conclusion aligns with previous reports describing stable heterochromatin domains induced by forced HP1 recruitment (Hathaway et al. 2012; Yelagandula et al. 2023).

To determine whether HP1-induced heterochromatin repression depends on SetDB1, we knocked out or knocked down SetDB1 in cells stably expressing the reporter and rTetR-HP1 transgenes. Cells lacking SetDB1 failed to repress the reporter upon HP1 recruitment (Fig. 1E), and SetDB1 knockdown significantly reduced the H3K9me3 signal on the reporter (Fig. 1F). These results indicate that SetDB1 is essential for HP1-induced heterochromatin establishment. Overall, the tethering experiments demonstrate that heterochromatin can be established through the recruitment of either SetDB1 (the writer of the heterochromatin mark) or HP1 (the reader). In the latter case, HP1 relies on SetDB1 to deposit the H3K9me3 mark, suggesting that heterochromatin formation involves a feedback loop rather than a simple linear pathway.

### HP1 directly interacts with SetDB1 by binding two histone H3-mimic motifs in unstructured regions of SetDB1

The dependence of HP1-induced silencing on SetDB1 suggests that HP1 may recruit SetDB1 through direct interaction. To explore this possibility, we expressed and purified both proteins. Flag-tagged human HP1 was purified from *E. coli* (Fig. S2), while EGFP-tagged SetDB1 was purified from HEK293T cells under high-salt (1M) condition to remove interacting proteins. Upon mixing the two independently purified proteins, SetDB1 and HP1 co-immunoprecipitated (Fig. 2A), indicating a likely direct interaction. Given the central roles of HP1 and SetDB1 in heterochromatin formation and function, we further characterized their interaction.

**Fig. 2.**
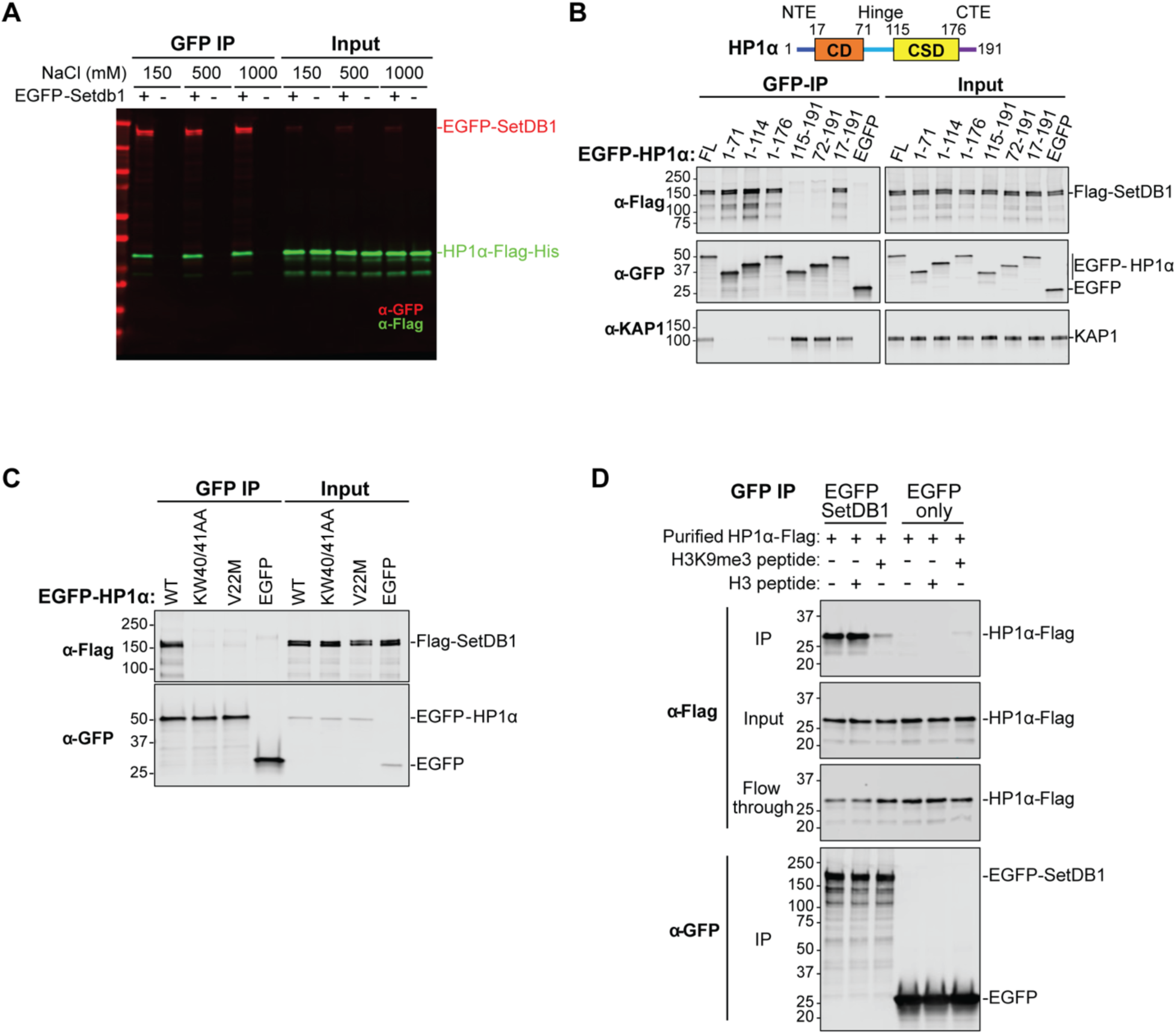
SetDB1 interacts with the chromodomain of HP1. (A) SetDB1 physically interacts with HP1α. EGFP-SetDB1, purified from HEK293T cells using wash conditions of different stringency (150 mM–1 M NaCl), was used to pull down HP1α-Flag-His purified from *E. coli*. Proteins were detected using anti-GFP and anti-Flag antibodies. (B) The chromodomain (CD), but not the chromoshadow (CSD) domain, of HP1α shows a strong interaction with SetDB1. A diagram of HP1α protein structure is shown at the top. Full-length (FL) or truncated fragments of EGFP-HP1α were transiently co-expressed with Flag-SetDB1 in HEK293T cells. Co-immunoprecipitation was performed using GFP nanotrap beads. KAP1, a known interactor of HP1α CSD, was detected for comparison. (C) HP1α mutants that disrupt the HP1-H3K9me3 interaction also impair HP1-SetDB1 interaction. WT or mutant EGFP-HP1α was transiently co-expressed with Flag-SetDB1 in HEK293T cells, followed by co-immunoprecipitation using GFP nanotrap beads and western blot analysis. (D) H3K9me3 peptide competes with SetDB1 for HP1α binding. EGFP-SetDB1 was purified from HEK293T cells using GFP nanotrap beads. HP1α-Flag-His purified from E. coli and synthesized histone peptides (methylated or unmethylated) were incubated with the bead-coupled EGFP-SetDB1. HP1α was detected in input, IP, and flowthrough fractions by western blot.

HP1 comprises two domains, the chromodomain (CD) and the chromoshadow domain (CSD), connected by an unstructured hinge region. The CD of HP1 forms a pocket that binds trimethylated lysine 9 on the histone H3 tail (H3K9me3) (Nielsen et al. 2002; Jacobs and Khorasanizadeh 2002), while the CSD mediates HP1 dimerization and interaction with multiple partners (Brasher 2000; Thiru et al. 2004; Hediger and Gasser 2006, 1). Many HP1-binding proteins possess a PxVxL motif that interacts with the CSD, and mutation of the conserved W174 residue in the CSD disrupts interactions with PxVxL-containing proteins (Brasher 2000; Thiru et al. 2004). To identify the interaction interface between SetDB1 and HP1, we generated truncated HP1 fragments and a W174A point mutant. Co-immunoprecipitation experiments revealed that HP1 fragments containing the CSD but lacking the CD failed to interact with SetDB1 (Fig. 2B). In contrast, KAP1, which possesses a PxVxL motif, interacted with the HP1 CSD as expected (Fig. 2B). Additionally, the W174A mutation did not affect HP1 binding to SetDB1 (Fig. S3), further suggesting that HP1 interacts with SetDB1 via a molecular interface distinct from that used for other partners. Notably, HP1 fragments containing the CD efficiently bound to SetDB1 (Fig. 2B).

The interaction between the HP1 chromodomain and trimethylated Lys9 on the histone H3 tail has been extensively characterized biochemically and structurally (Bannister et al. 2001; Lachner et al. 2001; Jacobs and Khorasanizadeh 2002; Nielsen et al. 2002). Several residues in the CD form conserved hydrophobic binding pockets for the histone tail carrying the K9me2/3 mark. Point mutations in residues V22 and W41 disrupted HP1 binding to the H3K9me3 peptide *in vitro* (Fig. S4). Surprisingly, the same mutations also disrupted HP1 interaction with SetDB1 (Fig. 2C), suggesting that HP1 uses the same molecular interface to bind both SetDB1 and the histone tail. If HP1 binds histone H3 and SetDB1 using the same pocket, these interactions would be mutually exclusive. Indeed, the addition of a trimethylated H3 peptide, but not an unmodified peptide, interfered with HP1-SetDB1 binding (Fig. 2D). These findings indicate that HP1 binds SetDB1 through its H3K9me2/3-binding pocket in the chromodomain.

We next sought to identify the interaction interface on SetDB1. Co-immunoprecipitation experiments with full-length HP1 and truncated SetDB1 fragments narrowed the interacting regions to two segments: amino acids 410–666 and 666–1291 (Fig. S5A). Using the AlphaFold2 Multimer algorithm (Jumper et al. 2021; Evans et al. 2022), we modeled potential complexes between full-length HP1 and these SetDB1 regions. AlphaFold predicted HP1 chromodomain binding to two motifs located in the unstructured regions of SetDB1 (Fig. 3A and Fig. S5B). Remarkably, these motifs resembled the H3 tail flanking Lys9, and their modeled 3D structures with HP1’s chromodomain were highly similar to the experimentally determined structure of HP1 bound to the histone tail (Fig. 3B and Fig. S5C) (Nielsen et al. 2002). The first motif, located in the ∼200-amino-acid unstructured region between the Triple Tudor and MBD domains, is highly conserved across mammals and insects (Fig. S6). The second motif, located within a ∼340-amino-acid unstructured region splitting the bifurcated SET domain, is conserved in vertebrates but absent in insects (Fig. S6).

**Fig. 3.**
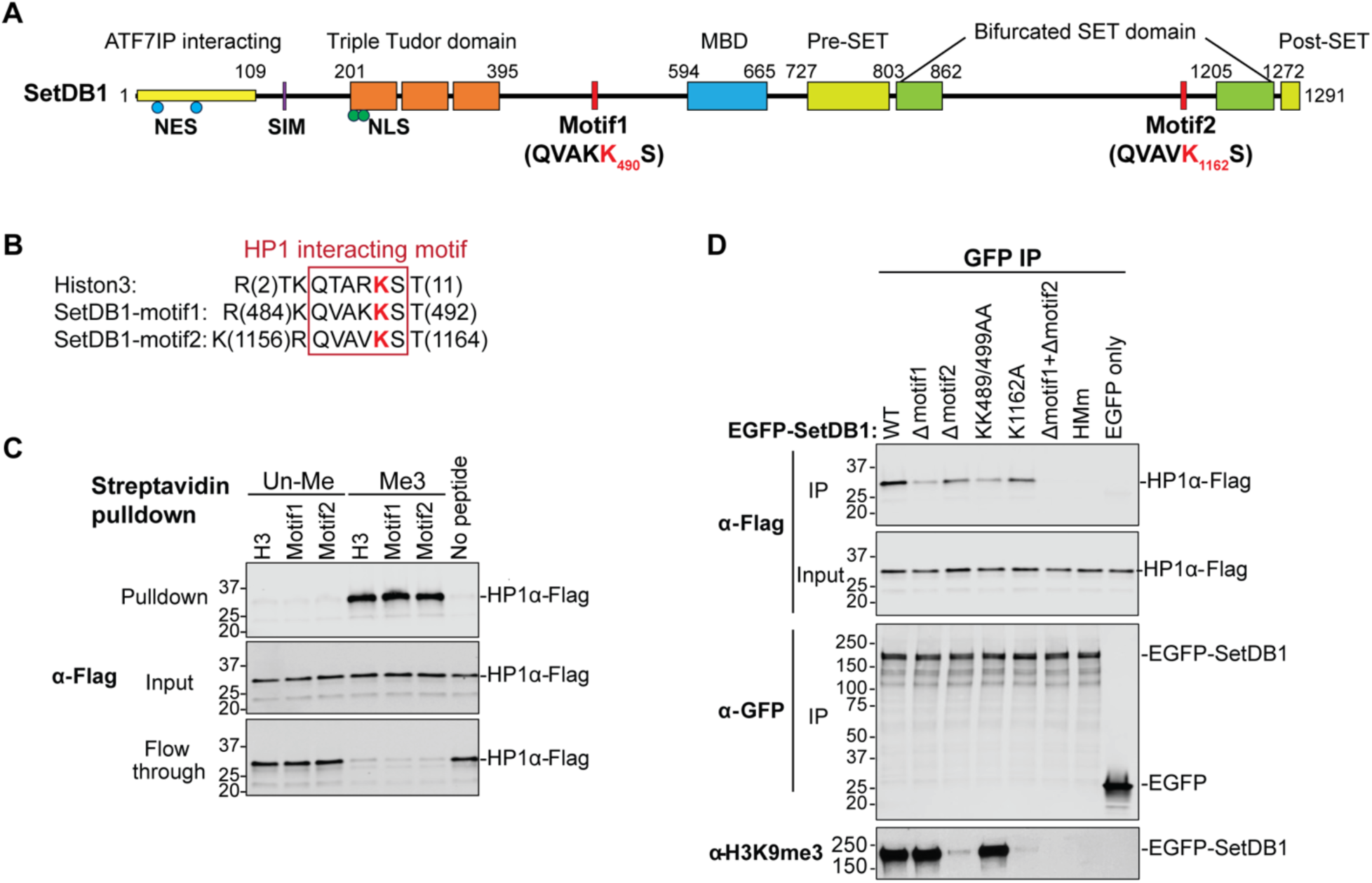
SetDB1 interacts with HP1 through two methylated histone mimic motifs (HMMs). (A) Domain architecture of SetDB1. Two HP1α-interacting motifs, predicted by AlphaFold2-Multimer, are highlighted, along with their corresponding sequences. (B) Sequence alignment of the histone H3 tail and the two identified SetDB1 motifs. H3K9 and the corresponding lysine residues in SetDB1 are shown in red. (C) HP1α interacts exclusively with methylated SetDB1 peptides *in vitro*. HP1α-Flag-His, purified from *E. coli*, was incubated with biotin-conjugated, lysine-trimethylated (Me3) or unmethylated (Un-Me) histone H3 or SetDB1 peptides. Pulldown was performed using streptavidin beads, and HP1α-Flag-His was detected by western blot. The negative control contained no peptide in the pulldown assay. (D) SetDB1 HMM mutants lose trimethylation and HP1 interaction. HP1α-Flag-His, purified from *E. coli*, was incubated with cell lysates from HEK293T cells expressing EGFP-SetDB1 (WT or mutants) and immunoprecipitated using GFP nanotrap beads. Cells expressing EGFP alone served as a negative control. HMm indicates a mutant with lysine-to-alanine mutations at positions 489, 490, and 1162. H3K9me3 antibody was used to detect trimethylated SetDB1.

To confirm HP1 binding to these motifs, we synthesized corresponding peptides and tested HP1 binding *in vitro*. Biotinylated peptides were incubated with purified HP1, followed by streptavidin pull-down and Western blot analysis. HP1 bound to both SetDB1 peptides with efficiency comparable to its binding to the H3 peptide (Fig. 3C). Importantly, HP1 binding occurred only when the Lys residues analogous to H3K9 in these peptides were trimethylated (Fig. 3C). These results demonstrate that HP1 interacts with two conserved histone H3-mimic motifs located in unstructured regions of SetDB1 in a methylation dependent manner.

To investigate the role of SetDB1’s histone-mimic motifs (HMMs) in HP1 binding in the context of the full protein, we generated SetDB1 mutants with either motif deleted, or Lys residues (corresponding to H3K9) mutated. For motif 1, in addition to mutating K490 (analogous to H3K9), we also mutated the upstream residue K489, as H3R8 has been implicated in HP1 binding (Jacobs and Khorasanizadeh 2002). Deletion or mutation of either motif reduced HP1 binding to SetDB1 (Fig. 3D). Simultaneous deletion or mutation of both motifs nearly abolished HP1 binding, demonstrating their necessity for efficient SetDB1-HP1 interaction. Overall, our findings reveal that the two conserved motifs in SetDB1 mimic the histone H3 tail and serve as binding platforms for HP1.

In addition to the three HP1 family members, several other chromodomain-containing proteins involved in heterochromatin repression, such as MPP8 (a component of the HUSH silencing complex), have been reported to bind the H3K9me3 mark (Chang et al. 2011; Tchasovnikarova et al. 2015). However, co-immunoprecipitation experiments showed that MPP8 binding to SetDB1 is substantially weaker than HP1 binding, indicating that not all chromodomain proteins interact efficiently with SetDB1 (Fig. S7).

### Histone-mimic motifs are autocatalytically modified by SetDB1 to create a complete mimic of the H3K9me3 mark

The pulldown experiments revealed that HP1 binds exclusively to trimethylated, but not unmodified, SetDB1 HMM peptides (Fig. 3C), suggesting that these motifs are modified in SetDB1 to provide an anchor for HP1 binding. To explore the modification of SetDB1 HMMs, we first used an antibody that specifically recognizes trimethylated lysine (K9) in the context of histone H3. We hypothesized that the similarity between the H3 tail and SetDB1 HMMs might allow this antibody to cross-react with SetDB1. Indeed, the anti-H3K9me3 antibody efficiently recognized SetDB1 purified from HEK293T cells (Fig. 3D). Deletion or point mutation of motif 2 strongly disrupted, but did not completely eliminate, antibody binding. Deletion of motif 1 had a milder effect on antibody recognition, while simultaneous disruption of both motifs completely abolished binding, indicating that both HMMs are modified by trimethylation (Fig. 3D).

To further confirm the modification of the HMMs, we conducted mass spectrometry analysis following thermolysin digestion of SetDB1 immunopurified from HEK293T cells. The mass spectrometry results revealed trimethylation of both HMM motifs, specifically K490 in motif 1 and K1162 in motif 2, corresponding to K9 in histone H3 (Fig. 4A). This provides additional evidence that the lysine residues corresponding to H3K9 in both HMMs are trimethylated *in vivo*.

**Fig. 4.**
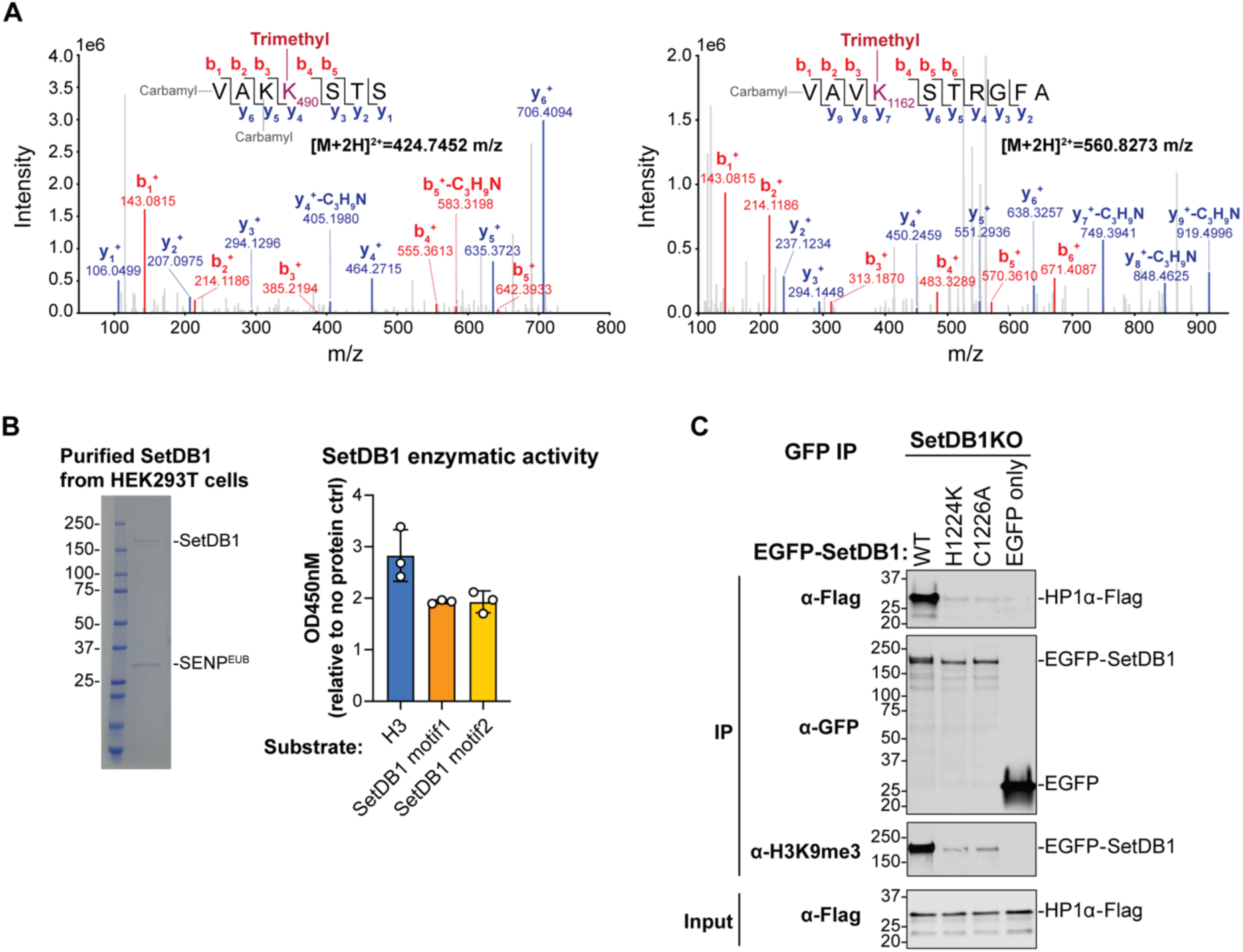
SetDB1 auto-methylates its histone mimic motifs *in vivo*. (A) SetDB1 K490 and K1162 can be tri-methylated *in vivo*. EGFP-SetDB1 was purified from HEK293T cells, digested with thermolysin, and analyzed by tandem mass spectrometry. Representative MS2 spectra of peptides with trimethylation at K490 (left, VAKKSTS, 424.7452 m/z) and K1162 (right, VAVKSTRGFA, 560.8273 m/z) are shown. The b and y ion peaks corresponding to the trimethylated peptides are labeled. The “-C3H9N” fragment indicates trimethylation. Carbamylation is an artificial modification occurring during thermolysin digestion. (B) SetDB1 methylates HMM motifs *in vitro*. EGFP-SUMO^EU^-SetDB1 was expressed in HEK293T cells, immunopurified using GFP nanotrap beads, and SetDB1 was eluted by SUMO protease SENP^EUB^ cleavage (left). Quantification of the methyltransferase activity of purified SetDB1 on histone H3 tail or SetDB1 HMM-containing peptides was performed using a commercial kit. Methylation of H3K9 was quantified using an HRP-conjugated secondary antibody-color development system at OD450 (right). Dots represent independent biological replicates; bars indicate mean ± SD. (C) SetDB1 lacking methyltransferase activity cannot efficiently interact with HP1α. EGFP-SetDB1-WT or methyltransferase-dead mutants were transiently expressed in SetDB1-knockout HEK293T cells. Cell lysates were incubated with HP1α-Flag-His purified from *E. coli*, followed by co-immunoprecipitation using GFP nanotrap beads and western blot analysis. EGFP alone was used as a negative control.

As SetDB1 is one of the two main enzymes in mammalian cells that deposits the H3K9me3 mark, we hypothesized that it might be responsible for modifying its own HMMs. To test this, we first explored whether SetDB1 modifies HMM peptides *in vitro*. SetDB1 purified from HEK293T cells methylated both HMM peptides in an *in vitr*o methylation assay (Fig. 4B). To determine whether SetDB1 methylates its own HMMs *in vivo*, we tested if impairing its methyltransferase activity affects HMM methylation. Previous studies have shown that point mutations in the catalytic SET domain, H1224K and C1226A, impair methyltransferase activity (Schultz 2002). We first validated these findings by expressing and purifying H1224K SetDB1 and demonstrating that its ability to modify H3 peptide was diminished in an *in vitro* methylation assay (Fig. S8). Next, we expressed these catalytically impaired SetDB1 mutants in HEK293T cells, which lacked SetDB1 due to CRISPR knockout of the endogenous SetDB1 gene. The significantly reduced recognition of both mutants by the anti-H3K9me3 antibody compared to wild-type SetDB1 and their inability to bind HP1 indicate a defect in HMM methylation (Fig. 4C). Thus, SetDB1 is the primary enzyme responsible for modifying its own HMMs. The residual modification observed in SetDB1 catalytically impaired mutants likely results from residual catalytic activity, as seen in the *in vitro* methylation assay (Fig. S8). Alternatively, another HMT, such as SUV39h, may contribute to the low level of methylation in the absence of SetDB1 activity.

Finally, we investigated whether SetDB1 performs intramolecular modification (*in cis*) or if one SetDB1 molecule methylates another *in trans*. To distinguish between these mechanisms, we purified the catalytically impaired SetDB1 mutants from cells expressing wild-type SetDB1. The presence of wild-type SetDB1 in these cells did not rescue the HMM methylation of the mutant SetDB1 (Fig. S9), indicating that the catalytic SET domain of SetDB1 preferentially acts on its own HMMs *in cis*. Together, these experiments reveal that SetDB1 is the primary enzyme responsible for its own intramolecular modification *in vivo*. More generally, they demonstrate that SetDB1 has non-histone substrates, extending its function beyond that of a writer of the histone code.

### The SetDB1 histone mimic motifs are required for heterochromatin spreading and maintenance

To explore the role of SetDB1 auto-methylation in SetDB1’s function, we first analyzed the catalytic activity of SetDB1 with disrupted HMM modification. In these and subsequent experiments, we used a SetDB1 mutant in which three lysine residues in the two HMMs were substituted (KK489/490AA and K1162A, referred to as HMm). These substitutions disrupt methylation of the two HMMs and prevent HP1 binding to SetDB1 (Fig. 3D). An *in vitro* methylation assay quantifying methyltransferase activity, using a histone H3 peptide as a substrate, showed that the methyltransferase activity of purified HMm SetDB1 was similar to that of the wild-type protein. This suggests that methylation of the HMMs is dispensable for SetDB1’s catalytic activity *in vitro* (Fig. S8).

Next, we investigated whether HMm SetDB1 could rescue SetDB1 deficiency in cells. Consistent with previous reports (Timms et al. 2016; Tsusaka et al. 2019), CRISPR-mediated knockout of the SetDB1 gene in HEK293 cells destabilized its cofactor ATF7IP, which is required for SetDB1’s nuclear localization and function (Fig. S10A). Stable expression of either wild-type or HMm SetDB1 restored ATF7IP expression, indicating that HMM methylation is not required for SetDB1-ATF7IP complex formation. Moreover, nuclear localization of SetDB1 was unaffected by the HMM mutation (Fig. S10B). Thus, disruption of HMM methylation does not impact SetDB1’s catalytic activity *in vitro,* nor its complex formation with ATF7IP or nuclear localization *in vivo*.

To further investigate the role of SetDB1 auto-methylation, we used the system for heterochromatin formation induced by the recruitment of chromatin factors, as described above (Fig. 1B). First, we examined the ability of HMm SetDB1 to induce transcriptional repression and heterochromatin formation upon direct recruitment to the reporter. A transgene stably expressing HMm SetDB1 fused to the rTetR DNA-binding domain was introduced into the cell line harboring the reporter, and recruitment was induced by the addition of Dox (Fig. 5A). Recruitment of HMm SetDB1 resulted in potent transcriptional repression, with efficiency and dynamics indistinguishable from those of the wild-type control. After 14 days of recruitment, >95% of cells with either wild-type or HMm SetDB1 exhibited silenced reporter expression (Fig. 5B). Recruitment of both the mutant and wild-type proteins led to similar levels of H3K9me3 and HP1 deposition at the TetO recruitment site (Fig. 5C, regions B and C), indicating that HMm SetDB1 retains normal catalytic activity toward chromatin *in vivo,* consistent with the *in vitro* findings (Fig. S8). However, heterochromatin spreading from the initial recruitment site was impaired. Specifically, levels of H3K9me3 and HP1 were reduced by more than 6- and 2-fold, respectively, on the hygromycin resistance gene located >5 kb upstream of the TetO site (Fig. 5C, region A). Consistent with ChIP-qPCR results, repression of the hygromycin resistance gene significantly decreased upon tethering of HMm SetDB1 (Fig. S11). Together, these results suggest that auto-methylation of SetDB1’s HMMs is dispensable for the establishment of heterochromatin around the initial recruitment site but is crucial for its spreading.

**Fig. 5.**
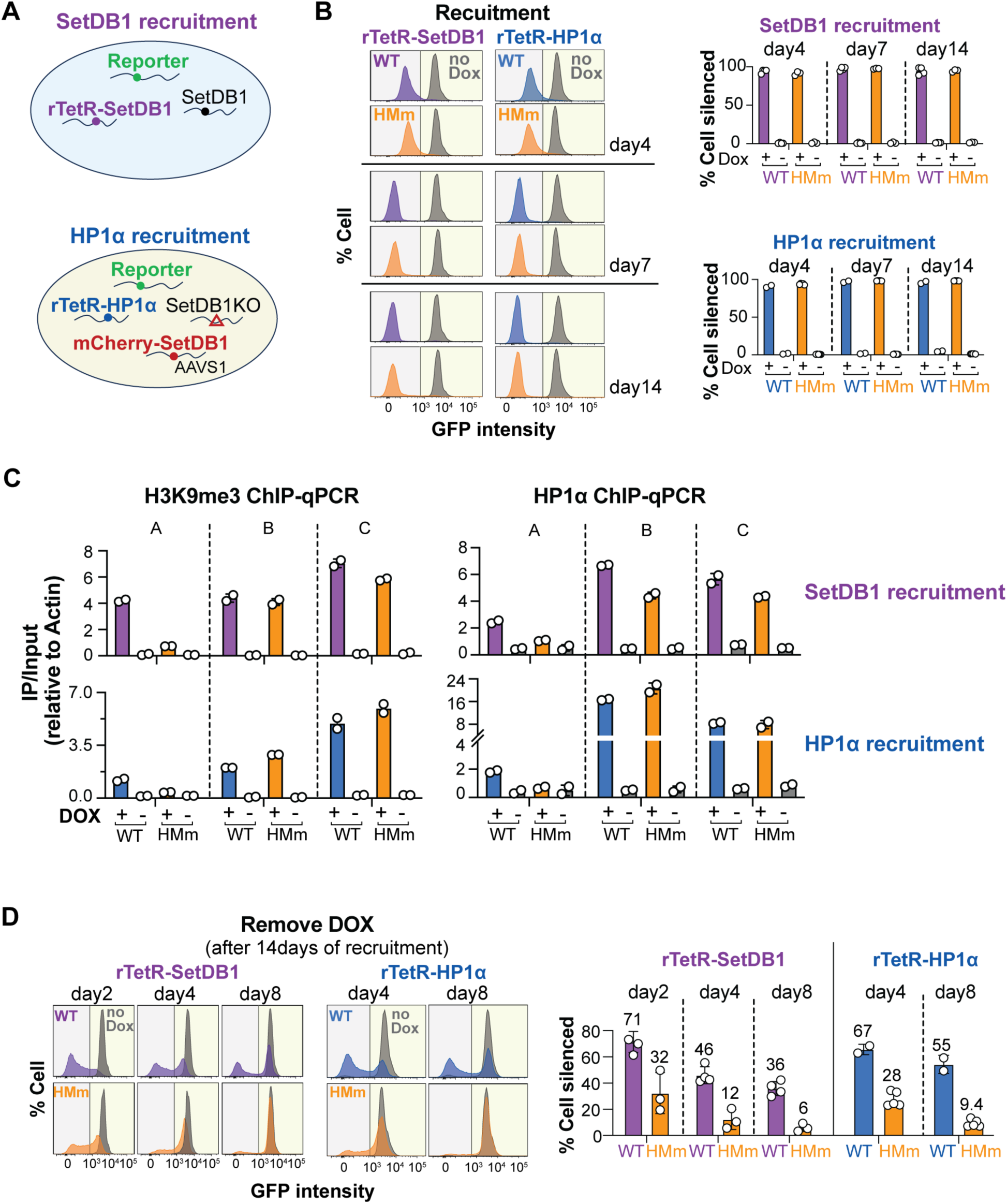
Loss of SetDB1 histone mimic methylation does not affect ectopic heterochromatin establishment on the reporter but impairs heterochromatin spreading and maintenance. (A) Schematic diagram of cells engineered to analyze the effects of HMM mutation (HMm) on SetDB1 (top) and HP1α (bottom) induced heterochromatin. For SetDB1 tethering, WT or HMm rTetR-SetDB1 was stably expressed in cells harboring the reporter. For HP1 tethering, WT or HMm mCherry-SetDB1 was stably expressed in SetDB1-KO cells expressing the reporter and rTetR-HP1α. (B) Reporter silencing upon HP1α and SetDB1 tethering is unaffected by SetDB1-HMM mutation. GFP signal was measured by flow cytometry upon SetDB1 or HP1α recruitment to the reporter over different periods (4, 7, and 14 days). The histograms (left) show results from representative single clones. The black line indicates the gating threshold to distinguish between GFP-positive (GFP+, yellow shading) and GFP-negative (GFP-, gray shading) cell populations. Dox-treated (purple/blue/orange) and no-Dox control (gray) samples are shown. Bar graphs (right) show the quantitative results from different clones. Dots represent single clones; bars indicate mean ± SD. (C) Local deposition of H3K9me3 and HP1α is unaffected by SetDB1-HMM mutation, but their spreading to distal regions is disrupted. H3K9me3 ChIP-qPCR (left) and HP1α ChIP-qPCR (right) were performed after 7 days of SetDB1 (top) or HP1α (bottom) recruitment. Dots represent independent biological replicates; bars indicate mean ± SD. See Fig. 1B for amplicon locations. (D) Defective SetDB1 histone mimic methylation impairs heterochromatin maintenance upon SetDB1 (purple) and HP1α (blue) recruitment. Dox was removed after 14 days of recruitment, and GFP signal was measured by flow cytometry at different time points after Dox removal. The histograms (left) show results from representative single clones. Bar graphs (right) show the quantitative results from different clones. Dots represent single clones; bars indicate mean ± SD. Numbers indicate mean values. Details and color scheme for histograms are the same as in panel B.

We next assessed whether SetDB1 auto-methylation influences heterochromatin formation induced by the recruitment of HP1. In the cell line harboring the reporter and rTetR-HP1 transgenes, we knocked out endogenous SetDB1 and integrated a transgene expressing either HMm or wild-type SetDB1 (Fig. 5A). HP1 recruitment triggered reporter repression in cells expressing HMm SetDB1, with repression efficiency and dynamics comparable to those in cells expressing the wild-type protein (Fig. 5B). However, as with direct SetDB1 recruitment, the spreading of repressive chromatin was disrupted in cells expressing HMm SetDB1 (Fig. 5C and Fig. S11). Overall, these results show that SetDB1 auto-methylation is dispensable for the initiation of heterochromatin formation but is critical for efficient spreading.

We further analyzed the role of SetDB1 auto-methylation in the maintenance of heterochromatin. As mentioned earlier, the reporter was fully repressed in >95% of cells after 14 days of recruitment of either wild-type or HMm SetDB1 (Fig. 5B). However, after recruitment was terminated, repression maintenance was significantly impaired in the HMM mutant. Indeed, when wild-type SetDB1 was tethered, the reporter remained repressed in a large fraction of cells even several days after Dox removal (46% and 36% of cells after 4 and 8 days, respectively). In contrast, upon HMm SetDB1 tethering, only 12% and 6% of cells maintained reporter silencing after 4 and 8 days, respectively (Fig. 5D). Similar results were observed when heterochromatin maintenance was assessed following HP1 tethering. After Dox removal, strong disruption of heterochromatin repression was seen in cells expressing HMm SetDB1. While >50% of cells expressing wild-type SetDB1 maintained repression after 8 days of Dox removal, fewer than 10% of cells expressing HMm SetDB1 showed repression (Fig. 5D). Thus, SetDB1 auto-methylation is essential for the maintenance of heterochromatin, regardless of whether heterochromatin formation was initially induced by HP1 or SetDB1 recruitment.

In summary, the tethering experiments revealed that auto-methylation of SetDB1 HMMs is crucial for the efficient spreading of heterochromatin during its establishment and for the stable maintenance of heterochromatin in the absence of the heterochromatin-inducing signal.

### SetDB1’s HMMs are required for proper genomic distribution of heterochromatin

To explore the function of SetDB1 auto-methylation on a genome-wide scale we analyzed heterochromatin in HEK293 cells in which endogenous SetDB1 was knocked out and which expressed a HMm (or control wild-type) SetDB1 transgene. Genome-wide profiling of the H3K9me3 mark by ChIP-seq showed a redistribution of heterochromatin domains in cells expressing the HMM mutant (Fig. 6A and B). Slight reduction of H3K9me3 level was observed genome-wide but was particularly prominent on the X chromosome (Fig 6A). On the other hand, elevation of H3K9me3 signal was detected on some regions, including euchromatic sites, where H3K9me3 signal was weak to moderate in the control cells, resulting in formation of ectopic heterochromatin domains (Fig 6. A, B and C). Indeed, analysis of the distribution of H3K9me3 using extended 1Mb-genomic windows revealed a strong (>1.5-fold) increase in H3K9me3 signal over 132 of the total 2853 windows (4.6%) (Fig. 6B). Regions with >1.5-fold elevated H3K9me3 signal were found on all chromosomes except Chr19 and ChrX (Fig. S12). These regions had lower gene density and much lower protein-coding gene density, compared to the average gene density in the rest of the genome (Fig. S13A). Genes within these windows were enriched in non-coding RNAs (40%) and pseudogenes (38%), which only account for 29% and 25% of annotated genes genome-wide, respectively (Fig. S13B). In contrast, only 10% of genes within these regions were protein-coding, whereas protein-coding gene make up more than 30% of the annotated genes genome-wide (Fig. S13B). Furthermore, the expression of genes located in these regions was significantly lower compared to the average from other regions (Fig. S13C), with most genes showing no (92.6%, RPKM=0) or low (7%, RPKM=0-1) expression in wild-type cells and less than 0.5% of genes showing expression with RPKM over 1, whereas more than 15% of genes located in the rest of the genome are expressed at RPKM>1 (Fig. S13C). Thus, regions that gained H3K9me3 in the HMM mutant were predominantly gene-poor and devoid of active expression, despite their euchromatic localization and relatively low H3K9me3 levels.

**Fig. 6.**
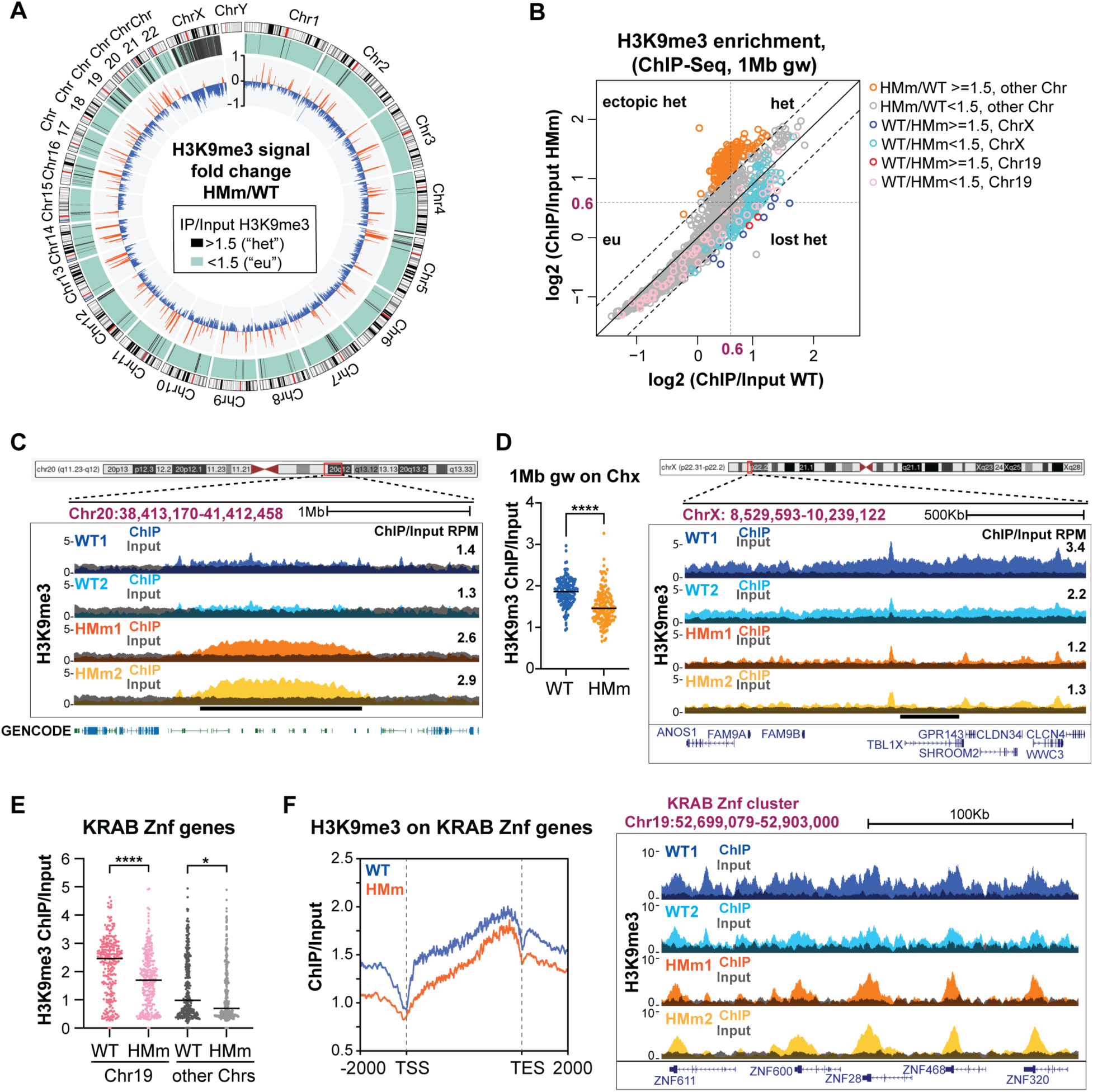
Loss of SetDB1 histone mimic methylation induces abnormal H3K9me3 distribution genome wide. (A) Genome-wide H3K9me3 distribution changes in SetDB1-HMM mutant. Circos plot showing H3K9me3 signal differences between cells stably expressing SetDB1-HMm or SetDB1-WT in a SetDB1-KO background. Outer gray tiles represent chromosomes. In the middle circle, black lines indicate 1Mb genomic windows where H3K9me3 ChIP/Input signal >1.5 in the WT rescue strain (defined as heterochromatic region, “het”). Green regions show no significant H3K9me3 enrichment (defined as euchromatin region, “eu”). The inner circle shows log2-transformed H3K9me3 signal changes (positive: orange, negative: blue) in SetDB1-HMm versus WT rescue strain. Data represent the average of two biological replicates. All subsequent ChIP-Seq and RNA-Seq data are from these two rescue strains. (B) The effect of SetDB1-HMm on H3K9 trimethylation varies across genomic regions. Scatterplot shows H3K9me3 ChIP/Input signals in 1Mb genomic windows (gw) from SetDB1-HMm versus WT rescue strain. Data represent the average of two biological replicates. Black dashed lines indicate a 1.5-fold difference. Gray dashed lines separate heterochromatic (H3K9me3 ChIP/Input >1.5) and euchromatic (H3K9me3 ChIP/Input <1.5) windows for both WT and HMm strains. Genomic windows from different chromosomes and with varying fold changes are depicted in different colors. (C) Euchromatin regions with low but detectable H3K9me3 signal gain H3K9me3 in SetDB1-HMM mutant. UCSC browser snapshot shows a representative euchromatin region with elevated H3K9me3. Tracks show overlaid ChIP (colored) and Input (gray) signals from two replicates. Numbers on the right show the normalized ChIP/Input signal for the manually selected genomic interval indicated by the black bar. (D) SetDB1-HMm leads to significant loss of H3K9me3 on ChrX. Left: Dot plot showing the distribution of H3K9me3 ChIP signal in 1Mb genomic windows from ChrX in both WT and SetDB1-HMm strains. Data represent the average of two biological replicates. The black line indicates the median. p<0.0001 is denoted as ****. Right: UCSC browser snapshot of a representative locus on ChrX. Tracks show overlaid ChIP (colored) and Input (gray) signals for two replicates. Numbers on the right show the normalized ChIP/Input signal for the manually selected genomic interval indicated by the black bar. (E) SetDB1-HMm leads to H3K9me3 loss over KRAB-Znf genes. Dot plot shows H3K9me3 signals on individual KRAB-Znf genes in SetDB1-WT and SetDB1-HMm strains. KRAB-Znf genes on Chr19 and on other chromosomes are plotted separately. Data represent the average of two biological replicates. The black line indicates the median. p<0.0001 is denoted as ****, p<0.05 is denoted as *. (F) H3K9me3 spreading on KRAB-Znf genes is impaired in SetDB1-HMM mutant. Left: Metaplot of H3K9me3 distributions over gene bodies of all KRAB-Znf genes in SetDB1-WT (blue) and HMm (orange) strains. Data is average of two biological replicates. Right: UCSC browser snapshot of a representative KRAB-Znf cluster on Chr19. Tracks show overlaid ChIP (colored) and Input (gray) signals for two replicates.

In contrast to genomic regions that acquired H3K9me3 signal in the SetDB1 mutant, genomic regions that lost H3K9me3 signal were non-randomly distributed. Indeed, H3K9me3 signal loss in SetDB1 mutant was particularly pronounced on chromosomes X and 19 (Fig. 6A and B, and Fig. S12 and Fig. S14). Of the 11 1Mb-genomic windows showing a strong (>1.5-fold) loss of H3K9me3 signal, 8 were located on ChrX and 2 on Chr19 (Fig. 6B). Around 50% and 20% of windows on ChrX and Chr19 showed >1.3x loss of H3K9me3 signal, respectively, while only 2% of windows on other chromosomes had the same level of H3K9me3 signal loss (Fig. S14). Further analysis revealed that in cells expressing the wild-type SetDB1, the X chromosome showed strong H3K9me3 enrichment compared to the other chromosomes (Fig. S15). This result is consistent with previous observations that in HEK293 cells, which are human embryonic kidney cells derived from a female with an atypical karyotype, two of the three X chromosomes are targeted for heterochromatin-mediated inactivation through the dosage compensation pathway (Gilbert et al. 2000). In cells expressing HMm SetDB1, H3K9me3 signal was lost along the whole X chromosome (Fig. 6D). Indeed, out of 151 1Mb-genomic windows on ChrX, 137 showed H3K9me3 loss (fold change >1) of which 77 showed over 1.3-fold loss and in 8 windows loss was over 1.5-fold. On a finer scale, although in SetDB1-HMM mutant heterochromatin decreased or was eliminated along most of the X chromosome, peaks of particularly high H3K9me3 enrichment remained mostly unaltered (Fig. 6D), suggesting a defect in spreading rather than initial heterochromatin formation. We employed RNA-seq to analyze if loss of heterochromatin affects the expression of genes on the X chromosome. Differential expression analysis revealed an increase in the expression of several genes located in heterochromatic ChrX windows which showed strong (>1.5-fold) loss of H3K9me3 signal in cells expressing HMm SetDB1 (Fig. S16A). Together, these results indicate that SetDB1 auto-methylation is involved in X chromosome inactivation possibly by enabling the spreading of heterochromatin from the initial entry sites.

Unlike the X chromosome, which shows H3K9me3 signal loss along its entire length in cells expressing HMm SetDB1, the H3K9me3 decrease on Chr19 occurs in large (500kb-2Mb) but distinct domains. These heterochromatin domains correspond to clusters of genes encoding KRAB zinc-finger (KRAB-ZNF) transcription repressors (Fig. S17A and B). Mammalian genomes encode hundreds of KRAB-ZNF proteins that recognize various transposable elements and endogenous retroviruses through DNA motifs. These proteins repress their targets by recruiting the KAP1 co-repressor and inducing H3K9 trimethylation via SetDB1 (Wolf et al. 2015; Schultz 2002). The expression of KRAB-ZNF genes is finely regulated through negative feedback, facilitated by KAP1-SetDB1-mediated establishment of heterochromatin over these genes (O’Geen et al. 2007; Frietze et al. 2010; Tchasovnikarova et al. 2015). We found that KRAB-ZNF genes located in clusters on Chr19, as well as those on other chromosomes, exhibit a loss of H3K9me3 in cells expressing HMm SetDB1 (Fig. 6E).

Examination of H3K9me3 profiles in wild-type cells over different individual KRAB-ZNF genes as well as metaplot analysis revealed that these genes exhibit the highest level of H3K9 trimethylation at their 3’ exons (Fig. 6F), confirming previous findings that heterochromatin is established on KRAB-ZNF genes by recognizing target sites located at their 3’ exons (O’Geen et al. 2007; Blahnik et al. 2011). Remarkably, while the strong enrichment of H3K9me3 over the 3’ exons of KRAB-ZNF genes was preserved in cells expressing HMm SetDB1, the H3K9me3 signal was reduced along the rest of the gene body and in flanking sequences (Fig6. F). These results suggest that SetDB1 auto-methylation is essential for the spreading of heterochromatin from the initial recruitment site at the 3’ ends of KRAB-ZNF genes, supporting the conclusions drawn from our tethering experiments (Fig. 5C). Additionally, we observed that the defective spreading of the H3K9me3 mark over KRAB-ZNF genes in HMm SetDB1-expressing cells led to a mild increase in gene expression (Fig. S16B). Overall, our findings demonstrate that SetDB1 auto-methylation is required for the spreading of heterochromatin and for proper repression of KRAB-ZNF genes.

## Discussion

### Histone mimic motifs facilitate complex formation of chromatin proteins

Histone proteins are subject to numerous post-translational modifications (PTMs), many of which act as chromatin marks that specify properties of distinct chromatin domains and regulate gene expression. Recent studies have identified that several non-histone proteins harbor short motifs resembling amino acid stretches present in histones. These histone mimic motifs (HMMs) can be post-translationally modified by enzymes that typically modify histones. However, whether HMM modifications are simply the result of off-target activity of the histone modifying enzymes or whether they have biological significance remains unclear. We discovered that the human histone methyltransferase (HMT) SetDB1 contains two histone mimic motifs. SetDB1 auto-modifies lysine residues within these motifs creating mimics of the H3K9me2/3 mark. The methylated HMMs are recognized by the chromodomain (CD) of HP1, enabling complex formation between SetDB1 and HP1. Our *in vivo* studies revealed that SetDB1 HMMs are critical for maintenance and spreading of heterochromatin providing direct evidence of HMMs playing a functional role in chromatin biology.

Compared to the large diversity of PTM in histones, HMMs identified to date in non-histone proteins are limited. A H3K4 mimic was reported in the influenza A virus nonstructural protein 1 (NS1) (Marazzi et al. 2012). The other experimentally verified HMMs, including those in SetDB1, mimic H3K9me2/3, the hallmark of heterochromatin. Notably, recipient proteins harboring H3K9 HMMs, such as SetDB1 and G9a, are themselves central to heterochromatin biology. G9a, in complex with GLP, catalyzes mono- and dimethylation of H3K9 and similar to SetDB1, auto-methylates its HMMs (Sampath et al. 2007; Chin et al. 2007). In addition, G9a modifies H3K9 HMMs in DNA Ligase 1 (Ferry et al. 2017) and in ATF7IP (Tsusaka et al. 2018), a co-factor of SetDB1 that is required for its nuclear localization and activity (Tsusaka et al. 2019; Wang et al. 2003).

The presence of H3K9me HMMs in SetDB1/ATF7IP and G9a/GLP complexes, which put this mark on histones, could stem from off-target activity of the HMTs. However, the evolutionary conservation of SetDB1 HMMs in otherwise poorly conserved unstructured regions suggests functional importance (Fig. S6). Proteome-wide analysis identified numerous putative HMMs and found their strong enrichment in chromatin-associated proteins (Chen et al. 2021), also implying functional relevance, rather than off-target proximity modifications.

What roles might HMMs play in SetDB1 and other chromatin proteins? One possibility is regulation of enzymatic activity. In fission yeast, the HMT Clr4 is regulated by auto-methylation, which exposes its substrate binding pocket and promotes an active conformation (Iglesias et al. 2018). Similarly, activity of the mammalian Suv39h2 was proposed to be regulated by auto-methylation, though with an opposite effect (Piao et al. 2016). However, our data show that auto-methylation of SetDB1 HMMs are not required for enzymatic activity. The *in vitro* enzymatic activity of the HMM mutant was indistinguishable from WT SetDB1’s and, importantly, the HMM mutant retained its ability to induce heterochromatin and reporter silencing *in vivo* (Fig. 5). Instead, SetDB1 HMMs mediate interaction with HP1, the key H3K9me2/3 reader (Bannister et al. 2001; Lachner et al. 2001). Other H3K9 HMMs also facilitate complex formation. The HMMs in G9a also interact with HP1 proteins (Sampath et al. 2007; Chin et al. 2007), while the HMM in DNA Ligase 1 recruits the DNA methyltransferase Dnmt1 to replication foci by interacting with the Tudor domain of UHRF1 protein (Ferry et al. 2017, 1; Kori et al. 2019) and the ATF7IP’s HMM mediates its interaction with MPP8 (Tsusaka et al. 2018), a CD-containing protein within the repressive HUSH complex. Thus, HMM modifications within SetDB1, G9a and ATF7IP enable their interactions with chromodomain proteins that otherwise act as readers of the H3K9me2/3 mark. Though *in vitro* binding studies or even co-immunoprecipitation from cellular extracts do not confirm physiological roles, they nevertheless suggest that HMMs expand the interaction network between histone marks, their readers and writers.

The discovery that HMMs in non-histone proteins are recognized by histone code readers adds another level of complexity to the biochemical interactions between chromatin components. Readers like HP1 and MPP8 bind H3K9me2/3 with distinct affinities (Chang et al. 2011). Their affinities towards the H3 tail and the HMMs in non-histone proteins likely also differ. Structural analysis of HP1 and HPP8 chromodomains in complex with H3K9me2/3 peptides show these interactions extend beyond the modified lysine (K9), involving flanking residues (Jacobs and Khorasanizadeh 2002; Nielsen et al. 2002; Chang et al. 2011). Differences in flanking sequences in histone H3 and HMMs of SetDB1 (Fig. 3B), G9a and ATF7IP, and the unique binding pockets of readers suggest varying binding affinities of diverse readers to HMMs and histone marks. In agreement with this, we observed SetDB1 HMMs interact more strongly with HP1 than with MPP8 (Fig. S7), whereas ATF7IP’s binding to MPP8 but not to HP1 seems to depend on its HMM (Tsusaka et al. 2018). Future studies should systematically assess reader affinities for HMMs and histone marks. It is plausible some readers prefer HMMs over genuine histone marks. Regardless, interactions between histone code readers and non-histone proteins modified by histone code writers broaden the concept of the histone code into a more general chromatin code.

### SetDB1 HMMs enable heterochromatin maintenance and spreading

Numerous studies have revealed complex regulation of heterochromatin, involving distinct stages of initial establishment, spreading and maintenance, each crucial for its role in nuclear organization and control of gene expression. Previous studies implicated HP1 as the main reader and identified SetDB1 and Suv39h1/2 as key writers of H3K9me3 (Padeken et al. 2022). Our results confirmed the central roles of HP1 and SetDB1 across all stages of heterochromatin function. We demonstrated that heterochromatin can be induced by transient recruitment of either SetDB1 or HP1 (Fig. 1 and Fig. S1). Since HP1 cannot deposit H3K9me2/3 by itself, it must cooperate with at least one HMT. Our data show that HP1-induced heterochromatin formation requires SetDB1 indicating that, at least in this experimental system, SetDB1 acts as the primary writer and Suv39h1/2 cannot compensate for its absence.

While SetDB1 is critical at all stages of heterochromatin formation, its HMMs are dispensable for initial heterochromatin establishment but essential for its spreading and maintenance (Fig. 5). This supports previous findings that establishment and maintenance stages are distinct and regulated by different mechanisms (Hall et al. 2002; Ragunathan et al. 2015; Tatarakis et al. 2023). Since SetDB1, but not its HMMs, is required for heterochromatin establishment upon HP1 tethering, there must be a mechanism enabling SetDB1 recruitment by HP1 independently of HMM binding. Indeed, coimmunoprecipitation experiments showed that SetDB1 binding to truncated HP1 lacking its chromodomain is strongly reduced but not eliminated (Fig. 2B). Many proteins bind HP1’s chromoshadow domain (CSD) dimer via PxVxL motifs (Thiru et al. 2004). A prior study and our experiments (not shown) revealed that HP1 CSD mutants defective in PxVxL interactions failed to induce heterochromatin formation upon tethering (Yan et al. 2018), suggesting these interactions recruit SetDB1. Recruitment could involve intermediaries like KAP1, which binds HP1 CSD via its PxVxL (Lechner et al. 2000) and SetDB1 in a SUMO dependent way (Ivanov et al. 2007). Future investigation into SetDB1 recruitment during heterochromatin establishment is needed to identify the molecular mechanism involved.

We found that SetDB1 HMMs are necessary for heterochromatin maintenance through cell divisions and for spreading along chromosomes – two critical features for heterochromatin function (Allshire and Madhani 2018; Grewal 2023). Reporter assays, in which establishment and maintenance can be studied separately, demonstrated that HMMs are dispensable for initial establishment but required for maintenance. The requirement of SetDB1 HMMs for spreading was apparent both in reporter experiments and in genome-wide analyses. SetDB1 HMM mutant showed reduced heterochromatin genome-wide, especially on the X chromosome (Fig. 6). In HEK293 cells, which have three ChrX copies, the entire X chromosome is heterochromatinized due to dosage compensation (Gilbert et al. 2000). H3K9me and SetDB1 play important roles in X inactivation (Keniry et al. 2016; Sun and Chadwick 2018; Minkovsky et al. 2014; Ichihara et al. 2022). In HMm-SetDB1 cells, most ChrX heterochromatin was decreased or lost, but prominent peaks remained unchanged, suggesting preserved heterochromatin nucleation but disrupted spreading.

X-chromosome inactivation is not the only instance exemplifying the importance of HMMs in heterochromatin spreading. Another affected region in SetDB1 HMM mutant is KRAB-ZNF gene clusters. SetDB1 recruitment to these clusters occurs via the KAP1-ZNF274 complex (and likely other KAP1-ZNF protein complexes) recognizing shared DNA motifs in the 3’ exons of KRAB-ZNF genes (Frietze et al. 2010), from where heterochromatin spreads across gene bodies and intergenic regions (Fig. 6). In HMM mutant, heterochromatin on KRAB-ZNF genes was reduced, yet the prominent H3K9me3 peaks at their 3’ ends were retained (Fig. 6).

Maintenance and spreading are vital for heterochromatin function. During development, heterochromatin represses lineage-specific genes (Nicetto et al. 2019; Wang et al. 2018; Ninova et al. 2019; Allshire and Madhani 2018; Padeken et al. 2022), with its maintenance ensuring sustained repression after initial silencing signals disappear. Spreading allows formation of extended heterochromatin domains, covering large genomic regions, such as pericentric heterochromatin, which supports 3D nuclear organization and repeat silencing (Maison and Almouzni 2004), or even entire chromosomes, such as during X inactivation (Loda et al. 2022). Extended transcriptionally silent heterochromatic domains are also formed in the euchromatic arms of chromosomes, such as at KRAB-ZNF (O’Geen et al. 2007; Barski et al. 2007) and odorant receptor gene clusters (Magklara et al. 2011). Finally, spreading is important on a smaller scale during the initial formation of local heterochromatin domains. Interestingly, KRAB-ZNF genes are not only targets of heterochromatin repression but also involved in heterochromatin formation over multiple copies of transposable elements (TEs) and endogenous retroviruses spread across the genome. KRAB-ZNF transcription repressors recognize DNA motifs within their targets and recruit the KAP1/SetDB1 silencing complex (Wolf et al. 2015; Imbeault et al. 2017; Ecco et al. 2016; Schultz 2002). Heterochromatin spreads from these recruitment sites, ensuring transcriptional repression of TEs. In germlines of both mammals and *Drosophila*, transcriptional silencing of TE is guided by non-coding piRNAs and this process also involves heterochromatin spreading to flanking sequences, including to adjacent genes (Pezic et al. 2014; Ninova et al. 2020; Thomas et al. 2013; Rozhkov et al. 2013; Sienski et al. 2012).

The simultaneous defects in maintenance and spreading in HMM mutant suggest a shared feed-forward mechanism sustaining heterochromatin by recruiting writers. In systems like *S. pombe* and the *Drosophila* germline, small non-coding RNAs (siRNA and piRNA) are produced from heterochromatin regions and guide writer recruitment to cognate genomic regions (refs). However, somatic mammalian cells lack small RNA-guided heterochromatin maintenance. An alternative ‘read-write’ mechanism could ensure a feed-forward loop if the same protein acts as both writer and reader of the heterochromatin mark. Indeed, *S. pombe* HMT Clr4 and the mammalian Suv39h1/2 contain N-terminal CDs, which *in vitro* were shown to bind to H3K9-methylated nucleosomes and methylate adjacent nucleosomes (Müller et al. 2016), suggesting a role in spreading and maintenance of heterochromatin. Clr4 mutant that diminished H3K9me binding impairs both heterochromatin spreading and maintenance (Zhang et al. 2008; Ragunathan et al. 2015). However, while SetDB1 plays a major role in heterochromatin function, a simple read-write mechanism cannot recruit SetDB1 as it can’t directly bind to H3K9me3 (Fig. S4, (Jurkowska et al. 2017)).

HP1 binding to SetDB1 HMMs suggests it might recruit SetDB1 to established heterochromatin. However, HP1 binds H3K9me3 and SetDB1 using the same CD pocket, preventing simultaneous binding (Fig. 2D). HP1 homodimerization through interaction of the CSDs might resolve this conflict as the two CDs within a dimer can bind two substrates. HP1 dimerization was suggested to stabilize condensed chromatin by binding H3 tails in two adjacent nucleosomes (Hiragami-Hamada et al. 2016; Azzaz et al. 2014). Alternatively, a dimer could bind H3K9me3 with one CD and a SetDB1 HMM with the other, thereby recruiting SetDB1 to heterochromatin (Fig. 7). This mechanism parallels direct read-write systems, enabling spreading and maintenance despite SetDB1’s inability to bind H3K9me3 directly.

**Fig. 7.**
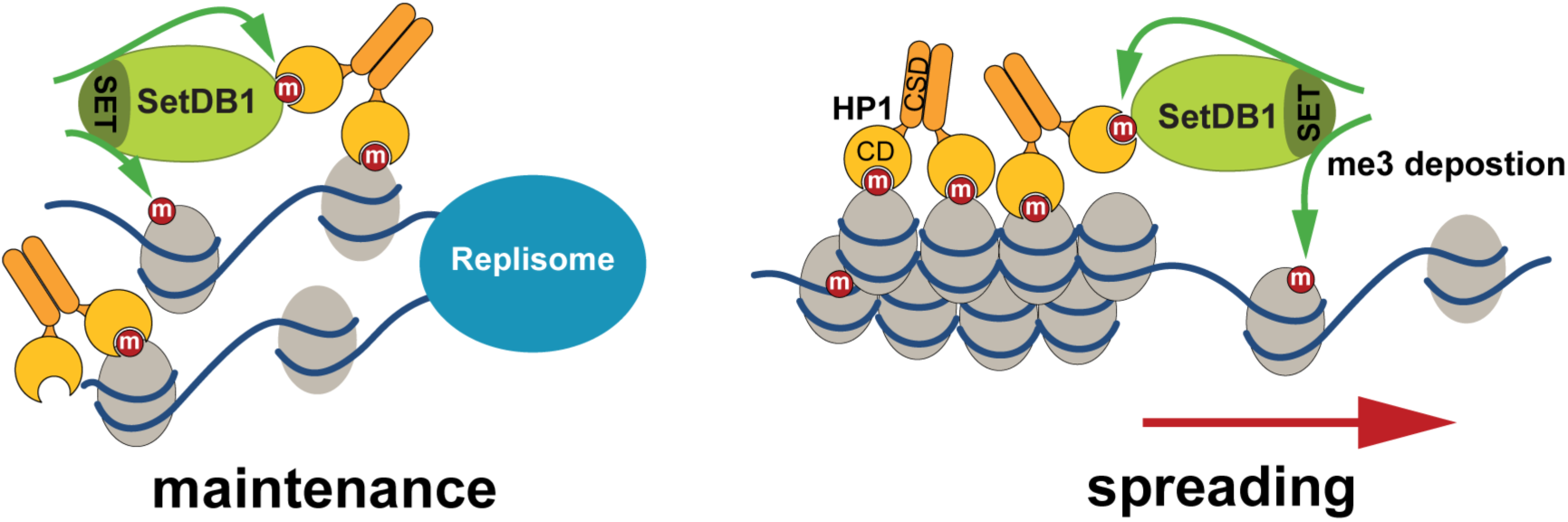
Model: Recruitment of SetDB1 to heterochromatin by HP1 enables heterochromatin maintenance and spreading. HP1 dimers were shown to stabilize condensed heterochromatin by binding H3K9me3 marks on adjacent nucleosomes. SetDB1 auto-methylates its HMMs creating sites for HP1 binding. In the indirect read-write model, SetDB1 is recruited to chromatin by HP1. In regions with lower density of the H3K9me3 mark, such as newly replicated chromatin (left) and the border region of heterochromatin (right), some HP1 dimers are not binding two histone tails, allowing one CD to bind and recruit SetDB1. SetDB1 methylates adjacent histone tails, maintaining heterochromatin integrity and enabling its spreading into adjacent regions. Accumulation of H3K9me3 displaces SetDB1 from chromatin as the H3K9me3 mark and SetDB1 compete for HP1 binding. Release of SetDB1 enables its redistribution to new, partially methylated regions.

Both direct and indirect HMT recruitment mechanisms may sustain heterochromatin through a feed-forward loop but the differences between the two mechanisms might explain the distinct functions and target preferences of SetDB1 and Suv39h. In mammals Suv39h1/2 predominantly maintains dense pericentric heterochromatin (Peters et al. 2001), while SetDB1 represses genes located at euchromatic arms of chromosomes, such as KRAB-ZNF genes, lineage-specific genes and endogenous retroviruses (Frietze et al. 2010; Tchasovnikarova et al. 2015; Bilodeau et al. 2009; Yuan et al. 2009; Karimi et al. 2011; Matsui et al. 2010; Geis and Goff 2020). Direct binding of Suv39h1/2 to H3K9me should result in recruitment that is proportional to H3K9me3 density, favoring H3K9me3-dense regions, such as their predominant target, pericentromeric heterochromatin. Indeed, a critical density of H3K9me3 is required for effective chromatin association of Clr4, the yeast homolog of mammalian Suv39h1/2 (Cutter DiPiazza et al. 2021). In contrast, SetDB1 may prefer regions with lower H3K9me3 density, allowing HP1 dimers to bind SetDB1 via free CDs. This might result in preferential SetDB1 recruitment to euchromatin-heterochromatin borders or regions where H3K9me3 was lost, e.g., due to replication, thereby promoting heterochromatin spreading and maintenance. In addition, competition with H3K9me3 for HP1 binding may evict SetDB1 from dense heterochromatin, allowing HP1 dimers to stabilize condensed chromatin and preventing ‘sponging’ of SetDB1 by dense heterochromatin and facilitating its relocation to yet-unmethylated regions. Thus, HMM-mediated SetDB1 recruitment might promote more efficient spreading and maintenance of heterochromatin compared to the direct read-write mechanisms (Fig. 7).

## Materials and methods

### Cell lines and cell culture

We used a commercial Flp-In-HEK293 cell line (Invitrogen, R75007), which contains a single stably integrated FRT site at a transcriptionally active genomic locus to generate stable cell lines. The FRT integrating locus was identified by inverse PCR (data not shown). To express proteins for coimmunoprecipitation, *in vitro* pull-down assays and *in vitro* histone methyltransferase reactions, we used HEK293T cells (gift from Varshavsky lab at Caltech). Cells were maintained in high glucose DMEM with GlutaMAX™ and sodium pyruvate (Gibco, 10569044) supplemented with 1% Penicillin-Streptomycin (10,000 U/mL, Gibco, 15140122) and 10% Tet Approved FBS (Takata, 631106) for stable cell lines for tethering assays, or 10% FBS (BenchMark, 100106) for other stable lines. 100μg/mL Zeocin (Invitrogen, R25001) was added to maintain the Flp-In-HEK293 cells. Cells were passaged at 80% confluency using 0.05% trypsin (Gibco, 25300054).

### Generation of the reporter cell line

The 5xTetO-pEF1α-EGFP-SV40pA reporter was cloned into the pcDNA5/FRT/TO vector backbone (Invitrogen V652020) by replacing the original pCMV-2xTetO_2_-MCS-BGHpA sequence. An intron sequence from an optimized splicing construct pPIP85.A (Moore and Sharp 1992) was placed in the middle of EGFP. 2x chicken HS4 insulators (Bintu et al. 2016) were placed upstream and downstream of the reporter to prevent effect from the surrounding genomic sequences. The EGFP fragment with intron was synthesized by gBlocks, the other elements of the reporter were amplified from a template plasmid (Addgene 78099). Integration of the reporter was performed by co-transfecting 200 ng of reporter plasmid and 1.8 μg of pOG44 plasmid (Invitrogen, V600520) into the Flp-In-HEK293 cells using Lipofectamine 3000 (Invitrogen, L3000008). pOG44 expresses a modified Flp recombinase, which mediates site specific integration of the reporter to the FRT site. Upon site-specific integration, expression of hygromycin resistance gene from the pcDNA5/FRT/TO vector was driven by a SV40 early promoter and a start codon located upstream of the FRT site in the Flp-In-HEK293 cell line. 48hrs post transfection, cells were transferred from 6-well plate to 10cm plate and selected with 100μg/mL hygromycin B (Invitrogen, 10687010) for 14 days. Single clones were isolated, and the integration of the reporter was verified by X-gal staining, Zeocin sensitivity test and genomic PCR, and a single clone was chosen for further experiments. Reporter line was maintained in DMEM supplemented with 100μg/mL hygromycin B.

### Integration of rTetR-HP1α or rTetR-SetDB1 into the reporter cell line

The rTetR fusion fragments (pEF1α-puro-T2A-rTetR-HP1α/SetDB1-BGHpA) were assembled into a vector backbone that contains PiggyBac inverted repeats. HP1α or SetDB1 fragments were synthesized by Twist bioscience, pEF1α was amplified from a template plasmid (Addgene, 78099), BGHpA was amplified from the pcDNA5/FRT/TO vector, and the puromycin resistance gene was amplified from the PiggyBac vector backbone. 1875 ng of rTetR fusion construct was transfected into the reporter cell line along with 625 ng of PiggyBac transposase plasmid using Lipofectamine 3000. Cells were selected with 1μg/mL puromycin (Gibco, A1113803) for 7 days, staring 48hrs post transfection. Single clones were obtained by limited dilution. Insertion of rTetR-HP1α (or SetDB1) was confirmed by western blot. Clones were maintained in DMEM medium supplemented with 100μg/mL hygromycin and 0.8μg/mL puromycin. At least two clones were chosen for the tethering assay.

### Generation of SetDB1 knockout cell lines

SetDB1 was knocked out using Crispr-Cas9 from two different cell lines, the rTetR-HP1α-reporter cell line and HEK293T cell line. To knock out SetDB1 from the rTetR-HP1α-reporter cells, two sgRNAs targeting SetDB1 exon2 (sgRNA1: GCATCCAAACCAATGCACCC) and exon3 (sgRNA2: CGTCCTCAGAGCTACTGTCC) were cloned into the pX333 vector (Addgene, 64073). 48hrs post transfection, cells were diluted and plated on a 96-well plate for single cell isolation. SetDB1 KO single clones were validated by genomic PCR, sequencing (Plasmidsaurus, Premium PCR) and Western blot. Two single clones were chosen for tethering assay, and one of them for further genome editing. To knock out SetDB1 from HEK293T cells, we cloned sgRNA1 into the pX330 vector backbone (Addgene, 42230) and constructed a donor plasmid for homology-directed repair by cloning a BSD-3xSTOP-SV40pA fragment flanked by SetDB1 homology arms (∼1000bp each) to the pJET1.2 vector backbone (Thermo Scientific, K1231). Upon recombination, expression of the blasticidin resistance gene was driven by the endogenous SetDB1 promoter. Cells were selected using 10μg/mL blasticidin (InvivoGen, ant-bl-05) and single clones were obtained by limited dilution and validated by genomic PCR and Western blot.

### Generation of SetDB1 rescue cell lines

To rescue the expression of WT and mutant SetDB1 in the SetDB1-KO cells, we integrated BSD-P2A-mCherry-SetDB1 (WT or mutant) at the first intron of the constitutively expressed *PPP1R12C* gene in the *AAVS1* locus of the rTetR-HP1α-reporter-SetDB1-KO cell line. We cloned the BSD-P2A-mCherry-SetDB1 (WT or mutant) fragment into an AAV zinc finger donor vector (Addgene, 22075) containing a promoter-less splice-acceptor and homology arms against the AAVS1 locus by replacing the puromycin resistance gene of the vector backbone. The integration was performed by co-transfecting 1µg of BSD-P2A-mCherry-SetDB1 donor plasmid and 500 ng of each TALEN arm targeting the AAVS1 locus (Addgene, 35431 targeting TGTCCCCTCCACCCCACA, and Addgene, 35432 targeting TTTCTGTCACCAATCCTG) into the rTetR-HP1α-reporter-SetDB1KO cell line. Cells were selected with 10μg/mL blasticidin (InvivoGen) and mCherry positive cells were sorted by fluorescence-activated cell sorting (FACS) with a Sony SY3200 cell sorter. Insertion of mCherry-SetDB1 at the AAVS1 locus was validated by genomic PCR and mCherry-SetDB1 expression was validated by western blot.

### Flow cytometry for reporter expression analysis

For artificial tethering in reporter cell lines stably expressing rTetR-HP1α (or SetDB1), media without hygromycin B and supplemented with Dox (1μg/mL) were used and changed every two days. To study maintenance of reporter silencing, Dox was completely removed from the media after extended tethering and cells were cultured in Dox free media for different periods of time. Cells without Dox treatment were cultured as “no Dox” control in parallel. Cells were harvested using 0.05% trypsin, resuspended in 0.22 μm-filtered flow cytometry buffer (Hank’s Balanced Salt Solution (Gibco, 14175079) with 2.5 mg/ml BSA (Sigma A4503)), and filtered through 35 μm strainers (Falcon 352235) to remove cell clumps. GFP fluorescence signal was measured with a MACSQuant VYB flow cytometer (Miltenyi Biotec) and a total minimum of 20,000 events were recorded for each sample. The data was analyzed with FlowJo and single cells were selected based on forward scatter Area vs. Hight (FSC-A vs. FSC-H). A manual gate was drawn on GFP fluorescence to determine the percentage of silent cells for each sample.

### HP1α expression and purification from *E. coli*

Codon optimized human HP1α-3xFlag was cloned into the pET28b (+) vector backbone and the plasmid was transformed into the LOBSTR *E. coli* strain. Expression was induced using 0.3mM IPTG at 18°C for 20 hrs. Cells were resuspended in lysis buffer (50 mM NaH_2_PO_4_, 600 mM NaCl, 20 mM imidazole, pH 8.0, cOmplete EDTA-free protease inhibitor cocktail (Roche 4693159001)) and lysed using a cell disruptor. Cell lysate was centrifuged at 10,000 g for 30 mins at 4°C. Clear lysate was incubated with Ni-NTA resin (Thermo Scientific, 88221) for 1hr at 4°C and beads were washed with ice-cold wash buffer (50 mM NaH_2_PO_4_, 600 mM NaCl, 50 mM imidazole, pH 8.0) and transferred to a bottom outlet capped column (BioRad, 7311550), followed by another two washes. Purified HP1α protein was eluted by elution buffer (50mM HEPES-KOH pH 7.5, 500mM NaCl, 250mM imidazole, 0.5mM TCEP) and dialyzed in buffer D (20 mM HEPES-KOH pH7.9, 0.2 mM EDTA, 100 mM KCl, 20% glycerol, 0.1mM PMSF, 0.5 mM DTT).

### Coimmunoprecipitation

EGFP and 3xFlag-fusion expression vectors were generated using the pcDNA3 vector backbone either one-by-one, or on the same vector separated by a P2A sequence. HEK293T cells were transfected with TransIT-LT1 (Mirus, MIR2304). Cells were harvested and lysed in cell lysis buffer (20mM Tris-Cl pH 7.5, 150mM NaCl, 1mM EDTA pH 8.0, 1% TritonX-100, cOmplete EDTA-free protease inhibitor cocktail) at 4°C, then lysate was cleared by centrifuging for 15 mins at the maximum speed (15000-20000xg) at 4°C. Cleared lysate was incubated with GFP-Trap magnetic agarose beads (Proteintech, gtma) for 2hrs at 4°C and beads were washed 3 times at 4°C with cell lysis buffer containing protease inhibitor and boiled in 2x Laemmli buffer (Sigma, S3401) for western blot analysis.

For detecting interaction between SetDB1 mutants and HP1α, pcDNA3-EGFP-SetDB1 plasmids were transfected into HEK293T cells and the cell lysate was incubated with 100nmol purified HP1α from LOBSTR *E. coli* and GFP-Trap magnetic agarose beads for 2hrs at 4°C, beads were washed and eluted in 2x Laemmli buffer.

### SetDB1 purification from HEK293T cells

EGFP-SetDB1 or EGFP-SUMO^EU^-SetDB1 were cloned into the pcDNA3 vector backbone and transiently transfected into HEK293T cells. Cells were lysed and incubated with GFP-Trap magnetic agarose beads as described above. EGFP-SetDB1 coupled beads were washed 3 times with cell lysis buffer containing 1M NaCl, then were washed and resuspended in cell lysis buffer with 150mM NaCl for pull-down assay. EGFP-SUMO^EU^-SetDB1 coupled beads were incubated with 250nM SUMO protease SENP^EUB^ ((Stevens et al. 2024), Gift from Voorhees lab at Caltech) in cleavage buffer (50mM Tris-Cl pH 7.5, 150mM NaCl, 0.1% TritonX-100, 1mM DTT) on ice for 1hr. The eluted SetDB1 was quantified and used for *in vitro* histone methyltransferase reactions.

### SetDB1 pull-down assay

For SetDB1 and HP1α interaction assay, EGFP-SetDB1 coupled beads were washed sequentially with high salt (1M NaCl) and normal (150mM NaCl) cell lysis buffer as described above and then were incubated with 100nmol HP1α in cell lysis buffer for 2hrs at 4°C. For the competition assay, the EGFP-SetDB1 coupled beads were incubated with 200μmol synthesized histone H3 peptides (with/without trimethylation on K9, Cayman Chemical, 10877 and 10530) and 10nmol HP1α in cell lysis buffer for 2hrs at 4°C. The beads were washed 3 times with cell lysis buffer and boiled in 2x Laemmli buffer. The supernatant was used for western blot analysis.

### Peptide pull-down assay

HEK293T cell lysates containing EGFP-fusion proteins or purified HP1α-3xFlag from LOBSTR *E. coli* were incubated with biotin conjugated histone H3 or SetDB1 HMM peptides and streptavidin beads (Invitrogen, 65001) in cell lysis buffer for 2hrs at 4°C. The beads were washed 3 times with cell lysis buffer and boiled in 2x Laemmli buffer. The supernatant was used for western blot analysis with GFP or Flag antibody. Biotin conjugated peptides were synthesized by GeneScript with over 90% purity (histone H3 (ARTKQTAR<u>K(with/without Me3)</u>STGGKAPRKQLA), SetDB1 HMM1 (QSRKQVAK<u>K(with/without Me3)</u>STSFRPGSVGSG), SetDB1 HMM2 (PMKRQVAV<u>K(with/without Me3)</u>STRGFALKSTHG).

### *In vitro* histone methyltransferase reactions

Quantification of the methyltransferase activity of purified human WT or mutant SetDB1 on histone H3 peptide and SetDB1 histone mimic motifs was performed with a commercial kit (Abcam, ab113453) according to the manufacturer’s instruction. The amount of peptides methylated by SetDB1 was quantified through HRP conjugated secondary antibody-color development system and color was read by a microplate reader (Tecan SPARK) at OD450 nm.

### ChIP-qPCR and ChIP-Seq

HEK293 cells were trypsinized and washed twice by 1xPBS (room temperature), then crosslinked using 1% formaldehyde for 10mins at room temperature. The reaction was quenched by glycine of a final concentration of 125mM and cells were washed twice with ice-cold 1xPBS and incubated in cell lysis buffer (50 mM Tris-HCl pH 7.5, 1mM EDTA pH 8.0, 140mM NaCl, 1% NP-40, cOmplete EDTA-free protease inhibitor cocktail) on ice for 10 mins. Chromatin was sheared by sonication in nuclear lysis buffer (50 mM Tris-HCl pH 7.5, 2 mM EDTA pH 8.0, 0.5% Sodium Deoxycholate, 0.35% SDS, cOmplete EDTA-free protease inhibitor cocktail) using a Bioruptor sonicator (Diagenode UCD-300) for 30 cycles (30s on/30s off) at high setting. The lysate was cleared by centrifugation at maximum speed (15000-20000xg) for 15 min at 4°C and diluted 3.5-fold in ice-cold dilution buffer (20 mM Tris-HCl pH 8.0, 1 mM EDTA pH 8.0, 0.1% Sodium Deoxycholate, 140 mM NaCl, 0.01% SDS, 1% NP-40, cOmplete EDTA-free protease inhibitor cocktail). The lysate was precleared by incubating with BSA blocked protein G beads (Invitrogen, 10004D) for 2-4hrs at 4°C. 5% of precleared lysate was saved as input, the rest was incubated with H3K9me3 or HP1α antibody overnight at 4°C, followed by incubation with blocked protein G beads at 4 °C for 4hrs. Beads were sequentially washed with low salt buffer (20 mM Tris-HCl pH 8.0, 2 mM EDTA pH 8.0, 0.1% SDS, 1% Triton X-100, 150 mM NaCl), high salt buffer (20 mM Tris-HCl pH 8.0, 2 mM EDTA pH 8.0, 0.1% SDS, 1% Triton X-100, 500 mM NaCl), LiCl buffer (10 mM Tris-HCl pH 8.0, 1 mM EDTA pH 8.0, 1% Sodium Deoxycholate, 1% NP-40, 250 mM LiCl) and TE buffer (10 mM Tris-HCl pH 8.0, 1 mM EDTA pH 8.0) at room temperature, followed by elution with elution buffer (50mM Tris-HCl, pH8.0, 10mM EDTA pH8.0, 1% SDS) at 65 °C for 1hr. Eluate and input were treated with RNase A and were reverse crosslinked at 65 °C overnight, then were treated by proteinase K at 55 °C for 2hrs. DNA was then extracted by phenol/chloroform. ChIP-qPCR was performed on a Mastercycler®ep realplex PCR thermal cycler machine (Eppendorf). Primers used in ChIP-qPCR are listed in Table1. All ChIPs were normalized to respective inputs and to the β-Actin control region. ChIP-seq library construction was carried out using the NEBNext® Ultra™ II DNA Library Prep Kit for Illumina (NEB, E7645). Libraries were sequenced using the Illumina NextSeq2000 system in SR100 mode.

### RNA extraction and RT-qPCR

HEK293 cells were washed twice with 1x PBS and RNA was extracted using TRIzol™ (Invitrogen, 15596018). RNA was treated with DNase I (Invitrogen, 18068015) and reverse transcribed using SuperScript III reverse transcriptase (Invitrogen, 18080085) with random hexamers primer (IDT). qPCR was performed on a Mastercycler®ep realplex PCR thermal cycler machine (Eppendorf) using primers listed in Table1.

### RNA-Seq

RNA was quantified using a Qubit RNA quantification kit (Invitrogen, Q32855) with a Qubit 2.0 fluorometer (Invitrogen). 1μg RNA was treated with DNase I (NEB, M0303) and depleted of ribosomal RNA with NEBNext® rRNA Depletion Kit (NEB, E7405). RNA-Seq libraries were made using the NEBNext® Ultra II Directional RNA Library Prep Kit (NEB, E7760). Libraries were sequenced on the Illumina NextSeq2000 system in SR100 mode.

### ChIP-Seq and RNA-Seq data analysis

The human genome assembly hg38 was used in all analysis. All alignments were performed through Galaxy (The Galaxy Community 2024), using Bowtie2 (Langmead and Salzberg 2012, 2) for ChIP-Seq datasets and STAR (Dobin et al. 2013) for RNA-Seq datasets.

ChIP-seq data was aligned allowing 3 mismatches and retaining only uniquely mapped reads. Genome coverage tracks were generated using deeptools-bamcoverage (Ramírez et al. 2016) through Galaxy and UCSC genome browser utilities ((Perez et al. 2024; Kent et al. 2010), normalized to the total numbers of uniquely mapped reads to the genome. For genome-wide H3K9me3 enrichment analysis, the genome was partitioned into 1Mb intervals. Interval coverage was calculated as the ratio of RPKM-normalized reads from ChIP libraries to Input libraries using deeptools-bamcoverage and custom R script. Low coverage intervals with fewer than 0.05 RPKM in Input libraries were excluded from the analysis. 1Mb genomic intervals with ChIP/Input ratio > 1.5 in WT ChIP-seq datasets (average value of two replicates) were annotated as “heterochromatin.” The full list of human KRAB-Znf genes was retrieved from the UniProt database and the H3K9me3 enrichment over individual KRAB-Znf genes was calculated as the ratio of the RPKM-normalized ChIP and Input read coverage within the annotated gene coordinates. The metaplot describing the average H3K9me3 enrichment (ChIP/Input) profile on KRAB-Znf gene bodies was generated using deeptools-computeMatrix and -plotProfile (Ramírez et al. 2016) through Galaxy. Circular plot was generated using Circos (Krzywinski et al. 2009) through Galaxy.

RNA-seq data was aligned allowing 3 mismatches and retaining only uniquely mapped reads. Reads number on annotated genes retrieved from Genecode was counted using featureCounts (Liao et al. 2014) through Galaxy. Differential expression analyses were performed using DESeq2 (Love et al. 2014) through Galaxy. For dotplots, read counts per gene were normalized as RPKM.

### Immunofluorescence (IF)

Cells were cultured on poly-D-lysine (Gibco, A3890401) treated chamber slides (Thermo Scientific, 177437PK), washed twice with 1xPBS, and fixed using 4% formaldehyde for 10 minutes at room temperature. Fixed cells were washed 3 times in 1xPBS and permeabilized with 0.5% TritonX-100 for 20mins at room temperature. Cells were blocked in blocking buffer (5% NGS and 0.3% TritonX-100 in 1xPBS) for 1hr at room temperature and then incubated with SetDB1 antibody (1:500, 3% NGS and 0.3% TritonX-100 in 1xPBS) overnight at 4°C. Cells were washed 3 times with 0.3% TritonX-100/PBS, followed by incubating with secondary antibody (1:500, 3% NGS and 0.3% TritonX-100 in 1xPBS) overnight at 4 °C in the dark. Cells were washed 3 times in 0.3% TritonX-100/PBS at room temperature in the dark and mounted with VECTASHIELD® PLUS Antifade Mounting Medium with DAPI (VectorLabs, H-2000-10). Imaging was performed using a Zeiss LSM 880 confocal microscope and data was processed using Fiji (Schindelin et al. 2012).

### SetDB1-HP1α interaction prediction by AlphaFold

AlphaFold2-Multimer (Jumper et al. 2021; Evans et al. 2022) was used to predict SetDB1-HP1α interactions on ColabFold (Mirdita et al. 2022). Full length HP1α was submitted to Colabfold along with aa 410-666 and aa 666-1291 of SetDB1, respectively. The predicted protein structures were analyzed using ChimeraX (Pettersen et al. 2021).

### Identification of SetDB1 methylation sites by mass spectrometry

EGFP-SetDB1 was expressed and purified from HEK293T cells using a modified cell lysis buffer (20mM Tris-Cl pH7.5, 150mM NaCl, 1% DDM, cOmplete EDTA-free protease inhibitor cocktail) and GFP-Trap magnetic agarose beads (Proteintech, gtma). EGFP-SetDB1 coupled beads were treated with TECP and CAA in a buffer containing 10mM HEPES pH 8.0 and 8M urea, followed by digestion with thermolysin (Promega) at 70°C with shaking for 3hrs in digestion buffer (40mM HEPES pH 8.0, 2M urea, 1mM CaCl_2_). Peptides were desalted using C18 Spin Columns (Pierce 89870) and eluted in elution buffer (70% ACN, 0.2% FA). All buffers and reagents were prepared using mass spec grade water.

LC–MS analysis of peptides was performed on an EASY-nLC 1200 (Thermo Fisher Scientific, San Jose, CA) coupled to a Q Exactive HF Orbitrap mass spectrometer (Thermo Fisher Scientific, Bremen, Germany) equipped with a Nanospray Flex ion source: 500 ng peptides were directly loaded onto an Aurora 25 cm × 75 μm ID, 1.6 μm C18 column (IonOpticks) heated to 50°C. The peptides were separated with a 72.5 min gradient at a flow rate of 350 nl/min as follows: 2%–6% Solvent B (3.5 min), 6%–25% B (42 min), 25%–40% B (14 min), 40%–98% B (1 min), and held at 98% B (12 min). Solvent A consisted of 97.8% H2O, 2% ACN, and 0.2% formic acid, and solvent B consisted of 19.8% H2O, 80% ACN, and 0.2% formic acid. The Q Exactive HF was operated in data-dependent mode with Tune (version 2.8 SP1 build 2806) instrument control software. Spray voltage was set to 1.6 kV, S-lens RF level at 50, and heated capillary at 275°C. Full scan resolution was set to 60,000 at m/z 200. Automatic gain control target was 3 x10^6^ with a maximum injection time of 15 ms. Mass range was set to 375–1500 m/z and charge state inclusion set to select precursors of charge state 2–5 for DDA analysis. For data-dependent ms2 scans, the loop count was 12, AGC target was set at 1×10^5^, intensity threshold was kept at 1×10^5^, and dynamic exclusion set to exclude precursors after one time for 45 seconds. Isolation width was set at 1.2 m/z and a fixed first mass of 100 was used. Normalized collision energy was set at 28. An inclusion list containing m/z values for predicted methylated peptides of SetDB1 was prioritized during data acquisition. Peptide match was set to off, and isotope exclusion was on. Data acquisition was controlled by Xcalibur (4.0.27.42), with ms1 data acquisition in profile mode and ms2 data acquisition in centroid mode.

Data analysis was performed using Thermo Proteome Discoverer 2.5 (Thermo Fisher Scientific, San Jose, CA) with a SEQUEST algorithm (PMID 24226387). The data was searched against a fasta file containing the sequence of SetDB1. The ms1 matching tolerance was 20 ppm, and the ms2 tolerance was 0.02 Da. Carbamidomethyl (+57.021 Da) on cysteine was set as static modification, and Oxidation (+15.995 Da) on methionine, and Carbamyl (+43.006 Da) on lysine were set as dynamic modification. Acetylation (+42.011 Da), Carbamyl (+43.006 Da), Met-loss (−131.040 Da), and Met-loss + Acetylation (−89.030 Da) at protein N-terminus were also set as dynamic modifications. A maximum of 2 miss-cleavages were allowed in the search. A concatenated target decoy-based percolator was utilized to control the false discovery rate. The q-value cutoff was set as 0.05. Protein abundances were reported using ms1 feature-based label-free quantitation. The median abundance for each sample was normalized to the same value.

### SetDB1 knockdown by siRNA

Silencer® Select siRNA against SetDB1 (ThermoFisher, 4427037) or Silencer Select control No. 1 siRNA (ThermoFisher, 4390843) was transfected into HEK293 cells with Lipofectamine™ RNAiMAX Transfection Reagent (Invitrogen, 13778030).

### Antibodies

Anti-H3K9me3 (Abcam, ab8898), anti-SetDB1 (Proteintech, 11231-1-AP), anti-FLAG (Sigma, F1804), rabbit polyclonal anti-GFP (Chen et al., 2016), anti-MPP8 (Proteintech, 16796-1-AP), anti-HP1α (Cell Signaling, 2616), anti-KAP1 (Invitrogen, MA1-2023), anti-ATF7IP (Proteintech, 14699-1-AP), anti-Tubulin (Sigma, T5168), IRDye® anti-rabbit and anti-mouse secondary antibodies (LICORbio, 926-32210, 926-32211, 926-68071, 926-68070), HRP-conjugated anti-rabbit and anti-mouse secondary antibodies (Cell Signaling, 7074 and 7076), Alexa Fluor 647-conjugated anti-rabbit secondary antibody (Invitrogen, A21245).

**Table1.**
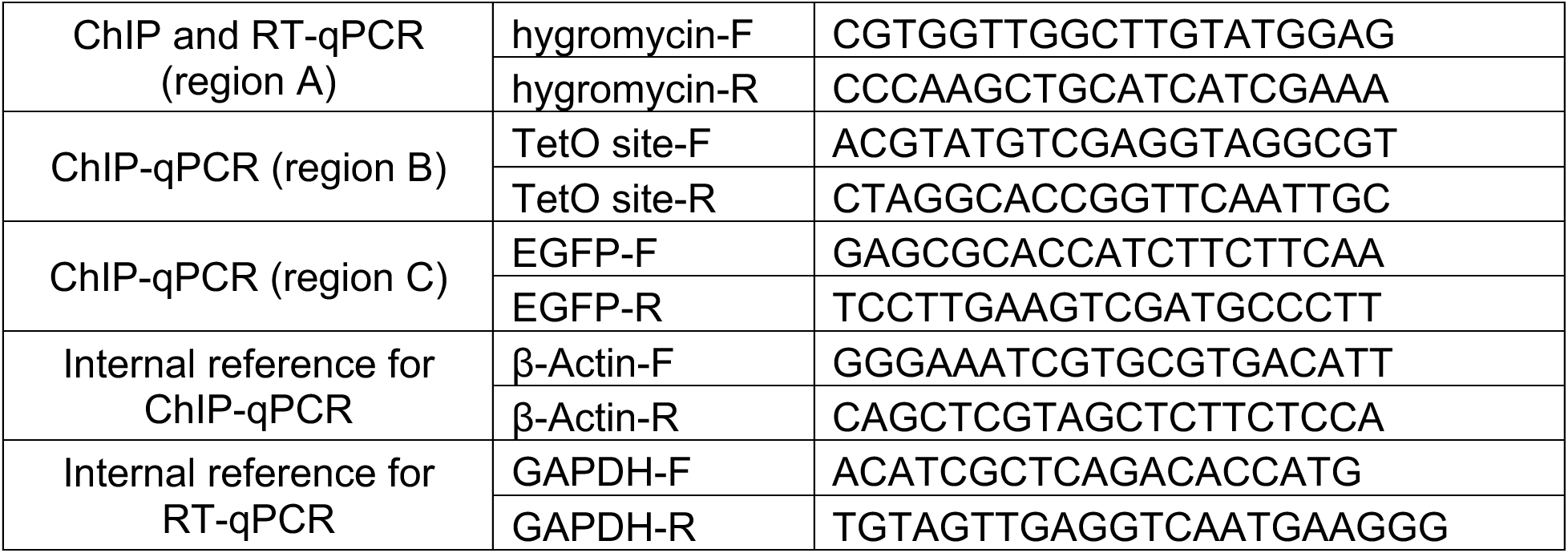
Primers for ChIP-qPCR and RT-qPCR.

## Acknowledgments

We thank members of the Fejes Tóth and Aravin labs for helpful discussions and comments. We are grateful to Akiko Kumagai and Masami Hazu for their suggestions and input on some of the experiments. Our thanks also go to Alexander Varshavsky for providing the HEK293T cell line, Michael Elowitz for sharing vectors, and Rebecca Voorhees for providing the SUMO protease SENP^EUB^. We appreciate the assistance of Jamie Tijerina, Diana Perez, and Rochelle Diamond with flow cytometry and FACS (Flow Cytometry and Cell Sorting Facility, Caltech); Baiyi Quan and Tsui-Fen Chou for their help with mass spectrometry (Proteome Exploration Laboratory, Caltech); Giada Spigolon and Andres Collazo for their support with imaging (Biological Imaging Facility, Caltech); Igor Antoshechkin for sequencing assistance (Millard and Muriel Jacobs Genetics and Genomics Laboratory, Caltech); and Maria Ninova for bioinformatics analysis. This work was supported by grants from the National Institutes of Health (R01 GM097363 to AA and R01 GM110217 to KFT) and by the HHMI Faculty Scholar Award to AAA.

## Author Contributions

Q.T., K.F.T. and A.A.A. conceived the study, designed the experiments, interpreted the results and wrote the manuscript. Q.T. performed experiments. M.S. helped with protein interaction experiments. Q.T. and A.Z performed the computational analysis and interpretation of the data. Q.T., K.F.T. and A.A.A. supervised the work, and K.F.T. and A.A.A. acquired funding.

---

**Fig. S1.**
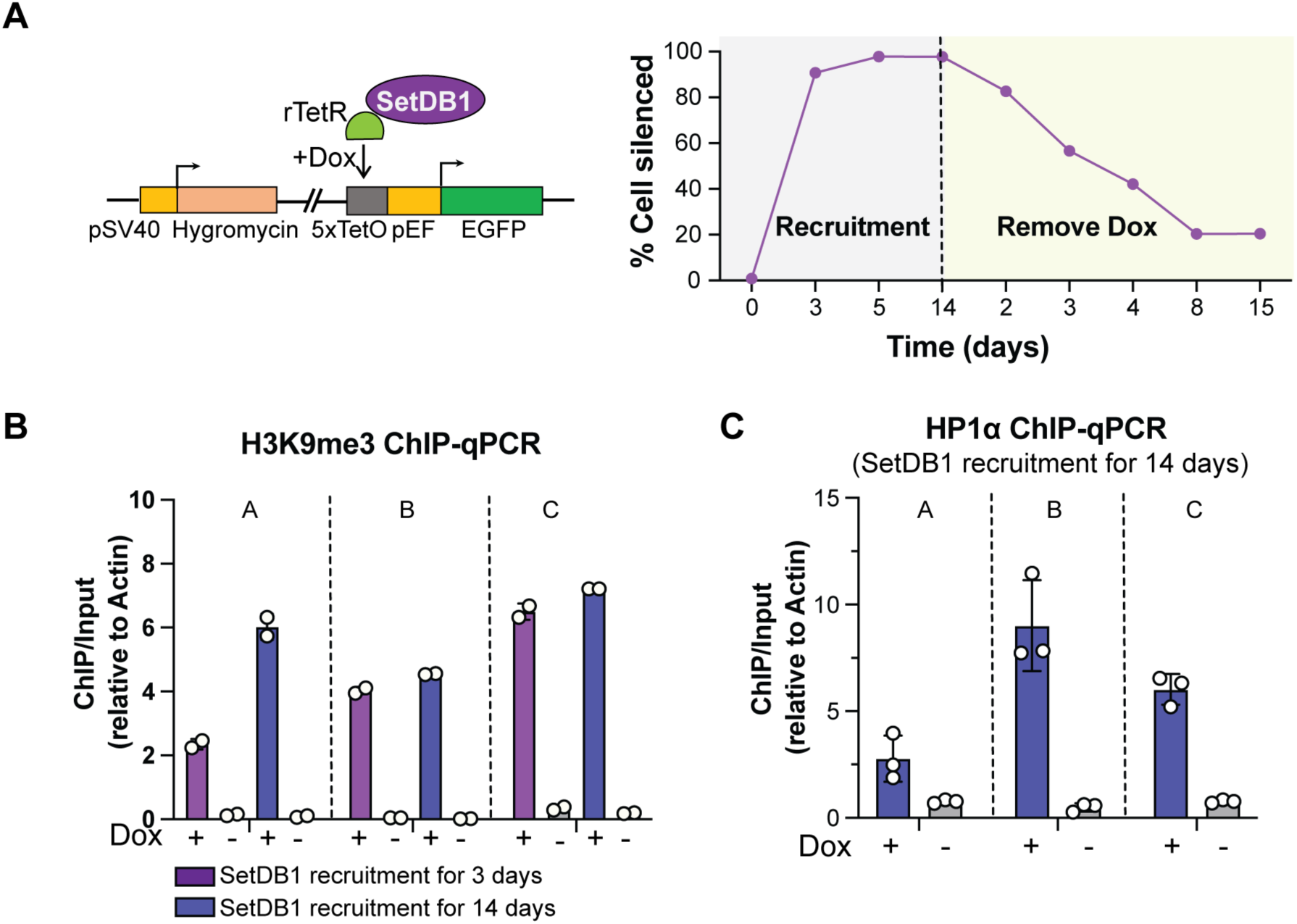
Recruitment of SetDB1 to the reporter leads to the formation of stable ectopic heterochromatin. (A) Artificial recruitment of SetDB1 leads to efficient silencing of the reporter. Left: Schematic diagram of the inducible reporter system. Right: rTetR-SetDB1 was recruited to the reporter upon Dox supplementation in the medium, and GFP signal was measured by flow cytometry at different time points. After 14 days of Dox treatment, Dox was removed from the medium, and flow cytometry measurements continued for an additional 15 days. Data represent the mean of 3 biological replicates. SD are plotted but are too small to be visible. (B-C) SetDB1 tethering induces *de novo* H3K9me3 and HP1 deposition along the reporter sequence. H3K9me3 ChIP-qPCR was performed after 3 and 14 days of SetDB1 recruitment (B). HP1α ChIP-qPCR was performed after 14 days of SetDB1 recruitment (C). The locations of amplicons along the reporter are indicated in Fig. 1B. Dots represent independent biological replicates; bars indicate the mean ± SD.

**Fig. S2.**
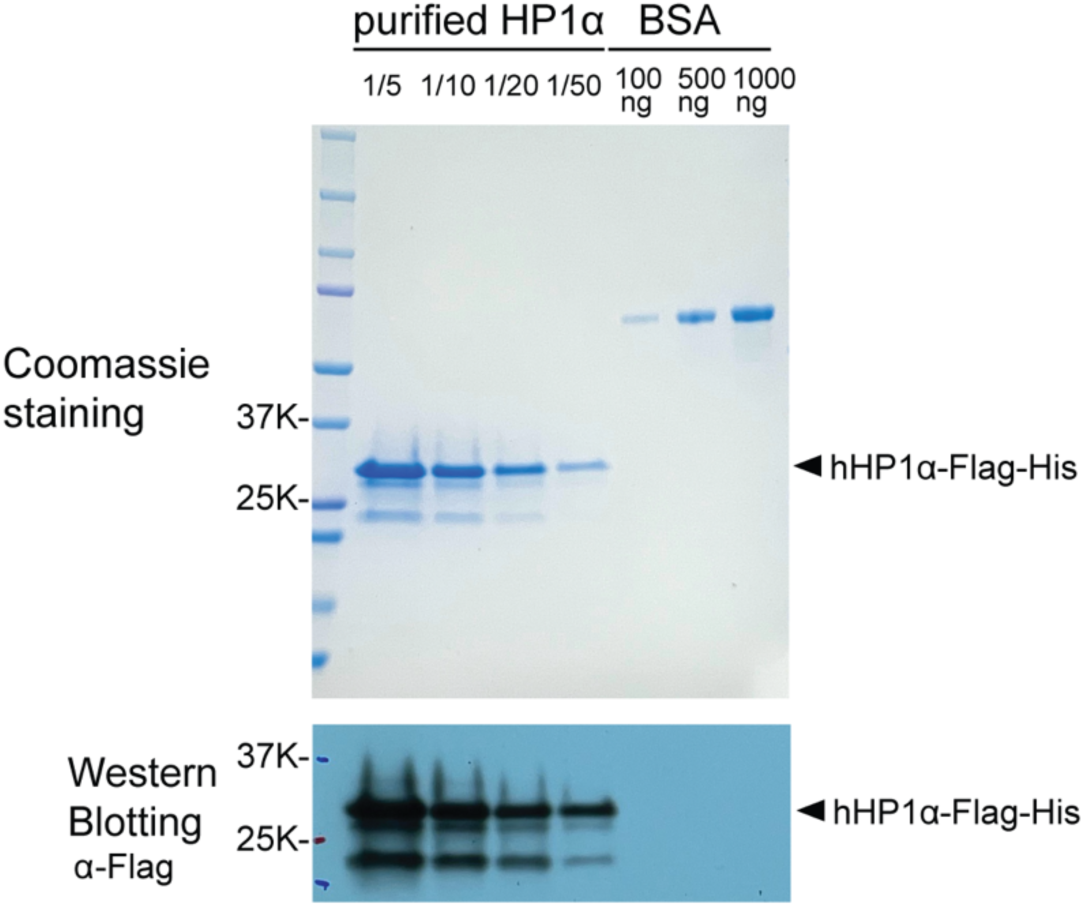
C-terminal Flag-His-tagged human HP1α was purified from *E. coli*. Coomassie staining and western blot were performed after affinity purification to assess the purity of the purified HP1α protein. Serial dilutions of purified HP1α and BSA were performed to determine the concentration of the purified HP1α protein.

**Fig. S3.**
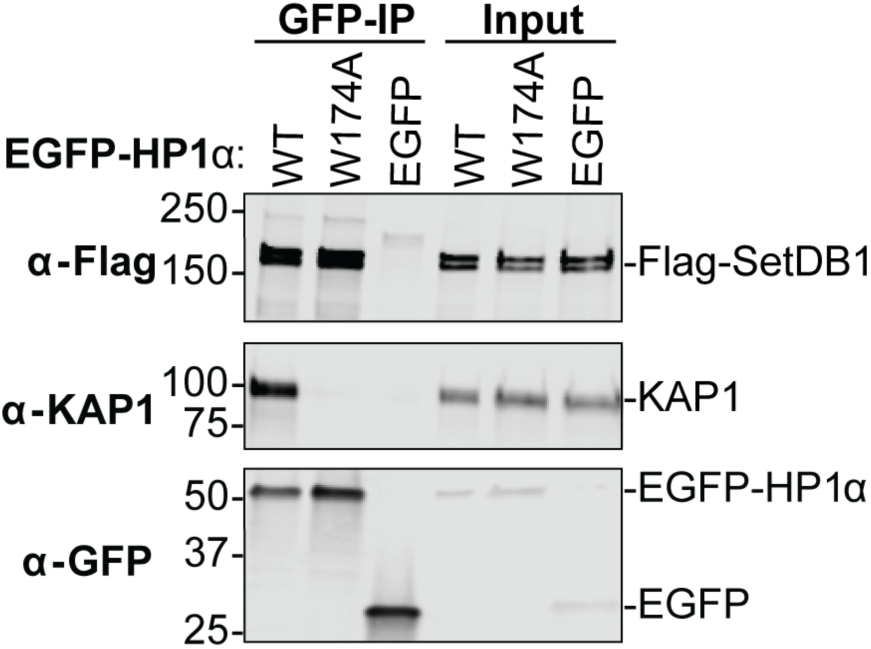
The W174A mutation of HP1α does not affect SetDB1-HP1 interaction. WT or W174A mutant EGFP-HP1α and Flag-SetDB1 were transiently co-expressed in HEK293T cells and co-immunoprecipitated using GFP nanotrap beads. The indicated proteins were detected by western blot. Cells co-expressing EGFP and Flag-SetDB1 were used as a negative control.

**Fig. S4.**
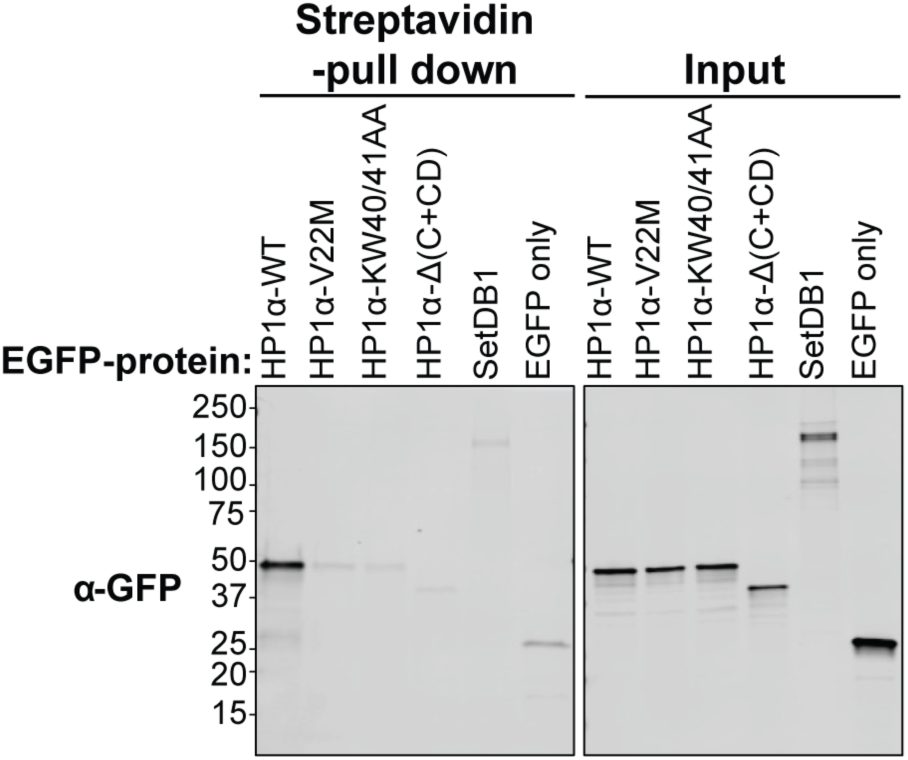
Mutations in the HP1α chromodomain disrupt HP1α-H3K9me3 interaction. Biotin-conjugated H3K9me3 peptide was incubated with cell lysate containing EGFP-HP1α and its respective mutants. Pulldown was performed using streptavidin beads, and EGFP-HP1α signal was detected by western blot. EGFP and EGFP-SetDB1 were used as negative controls.

**Fig. S5.**
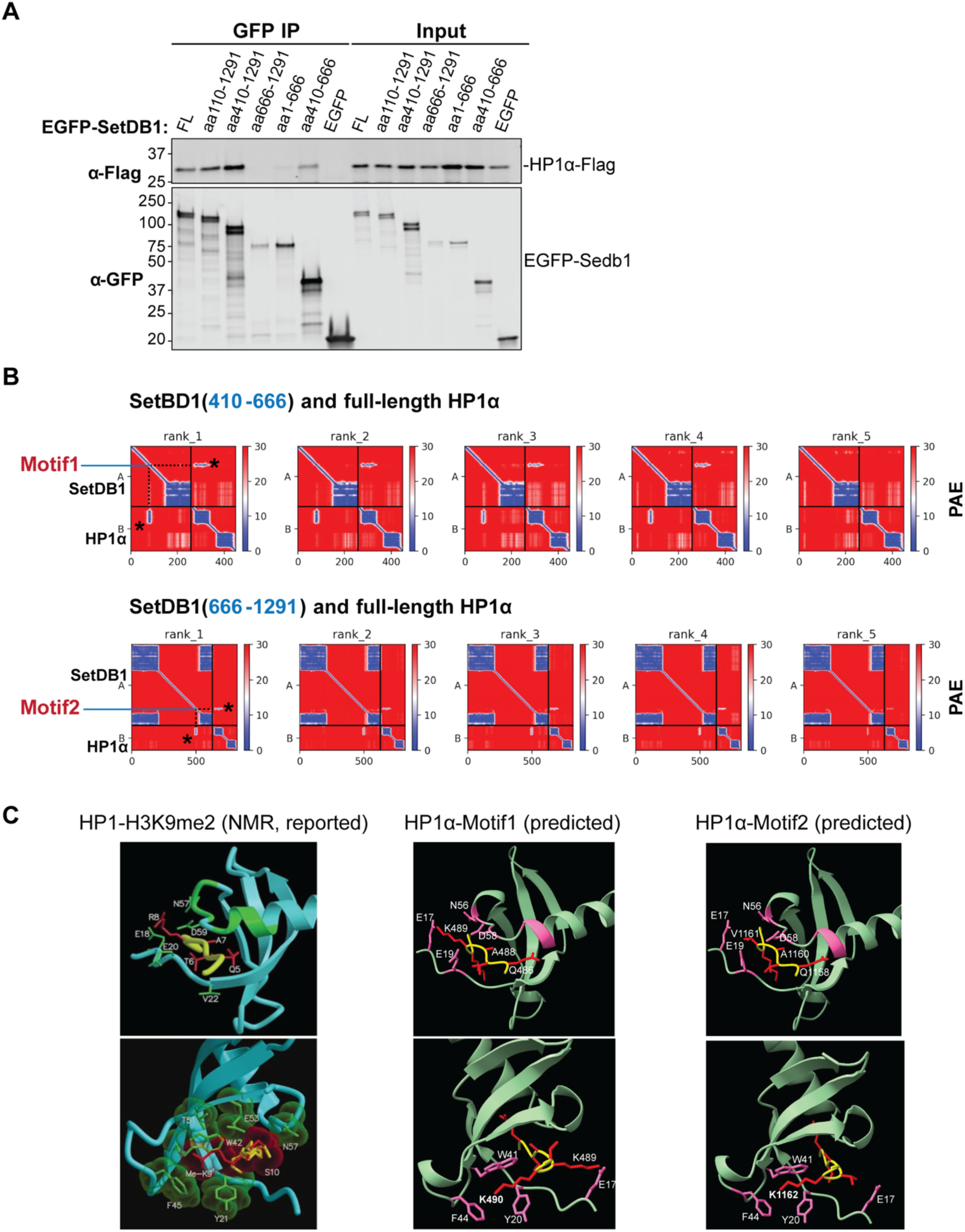
AlphaFold2 predicts two motifs in SetDB1 that interact with the HP1α chromodomain. (A) Two different regions (aa 410-666 and aa 666-1291) are predicted to interact with HP1α. Full-length (FL) or truncated EGFP-SetDB1 and HP1α-Flag were co-expressed in HEK293T cells. Co-immunoprecipitation was performed using GFP nanotrap beads, and western blot was carried out to detect the immunoprecipitating proteins. Numbers indicate the expressed amino acid stretches. (B) AlphaFold2 predicts two motifs on SetDB1 that interact with the HP1α chromodomain. Full-length HP1α was submitted to AlphaFold2-Multimer along with two different SetDB1 regions (aa 410-666 and aa 666-1291). The PAE (predicted aligned error) files show five different interaction models generated by AlphaFold2-Multimer for each prediction. The two SetDB1 regions with the lowest PAE values are labeled as motif 1 and motif 2. (C) The predicted structures formed by HP1α-SetDB1 interactions show high similarity to the reported HP1-H3K9me2 interaction. The HP1 chromodomain is shown in blue, with relevant residues forming the binding pocket in green (left, reported HP1-H3K9me2 interaction (Nielsen et al. 2002)) or in green with relevant residues marked in pink (middle and right, predicted HP1α-SetDB1 interactions). Histone H3, SetDB1 motif 1, and motif 2 are shown in yellow (main chain) and red (side chains).

**Fig. S6.**
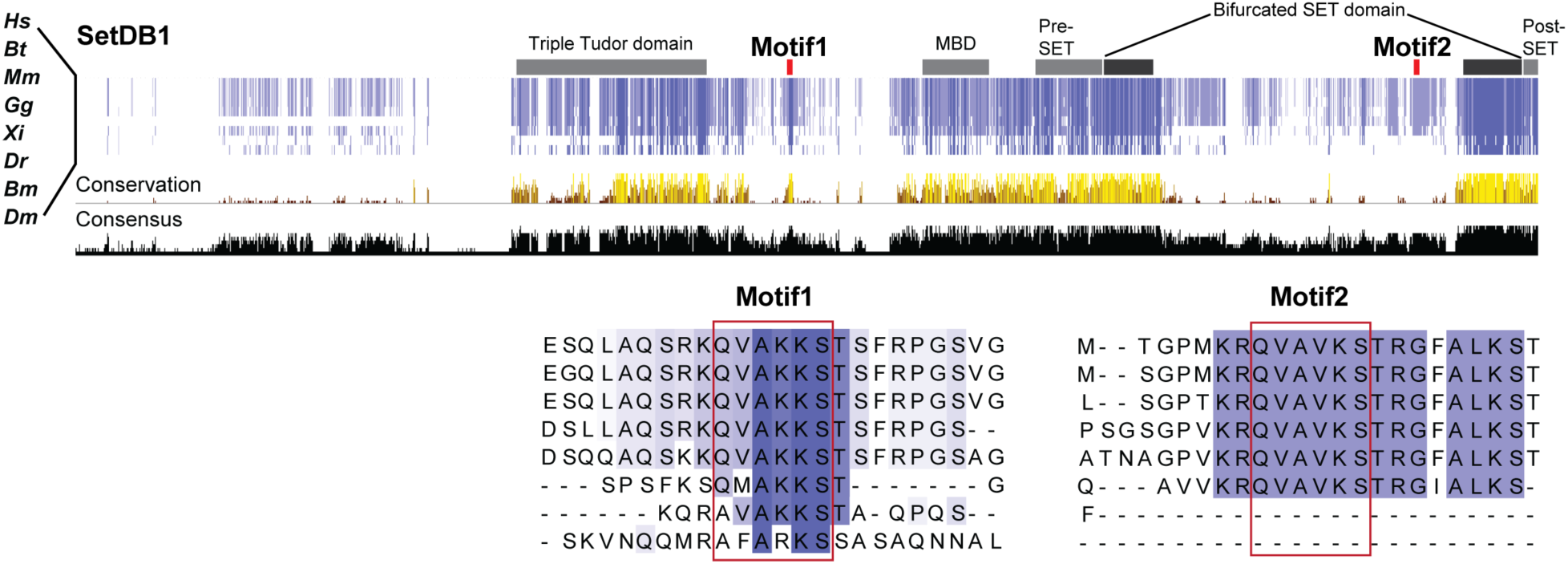
SetDB1 motif 1 is conserved from fly to human, and motif 2 is highly conserved in vertebrates. Sequence alignments of SetDB1 in different species (*Homo sapiens (Hs), Bos taurus (Bt), Mus musculus (Mm), Gallus gallus (Gg), Xenopus laevis (Xi), Danio rerio (Dr), Bombyx mori (Bm), Drosophila melanogaster (Dm)*) were performed using NCBI-COBALT. A schematic of positions colored by percentage identity is shown. Histograms reflect conservation and consensus. SetDB1 motif 1 and motif 2 are marked by red bars. The regions containing motif 1 and motif 2 and their flanking sequences are magnified (bottom). Images are based on alignments edited in Jalview (Clamp et al. 2004).

**Fig. S7.**
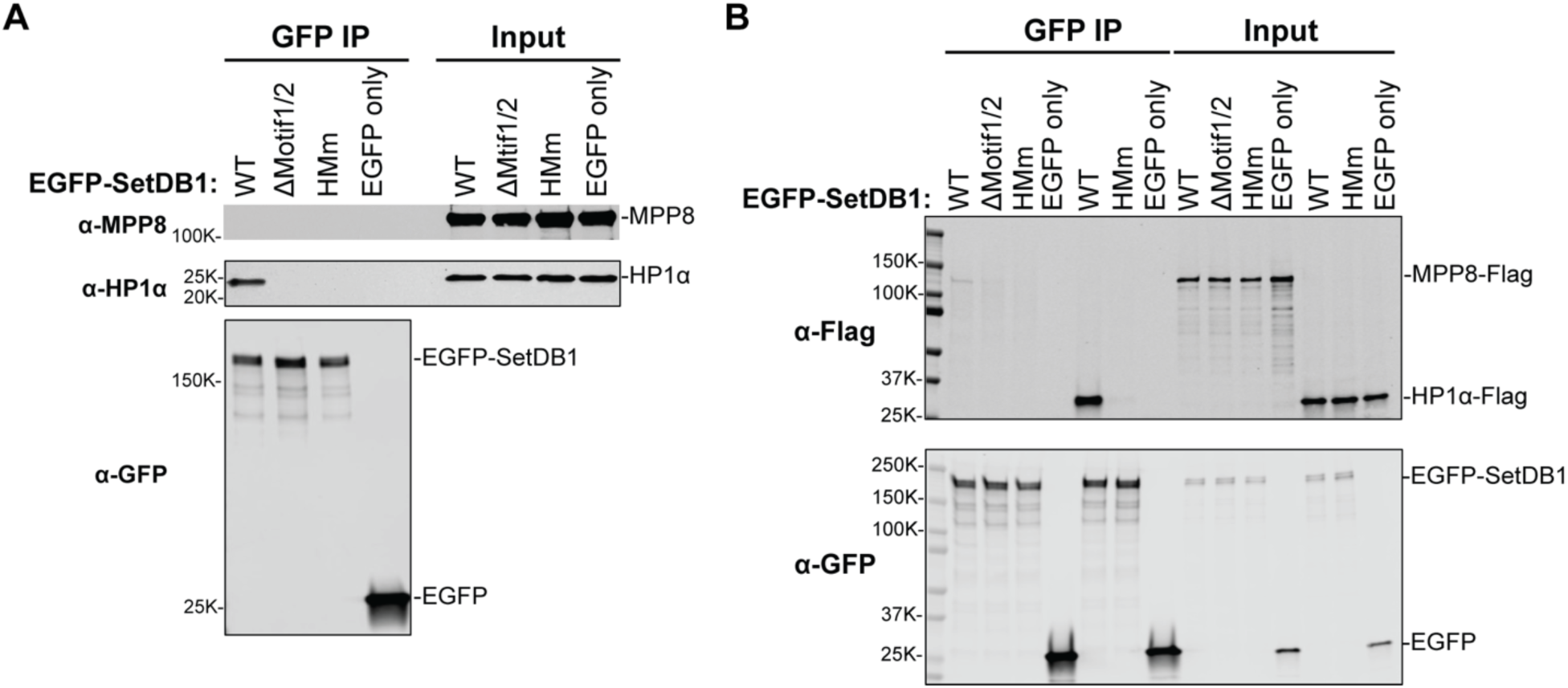
SetDB1’s interaction with MPP8 is much weaker than with HP1. (A) EGFP-SetDB1 and mutants were expressed and immunoprecipitated from HEK293T cells. Endogenous HP1α and MPP8 were detected by western blot. (B) EGFP-SetDB1 was co-expressed with MPP8-Flag or HP1α-Flag in HEK293T cells and immunoprecipitated using GFP nanotrap beads. Overexpressed HP1α and MPP8 were detected by Western blot using an anti-Flag antibody.

**Fig. S8.**
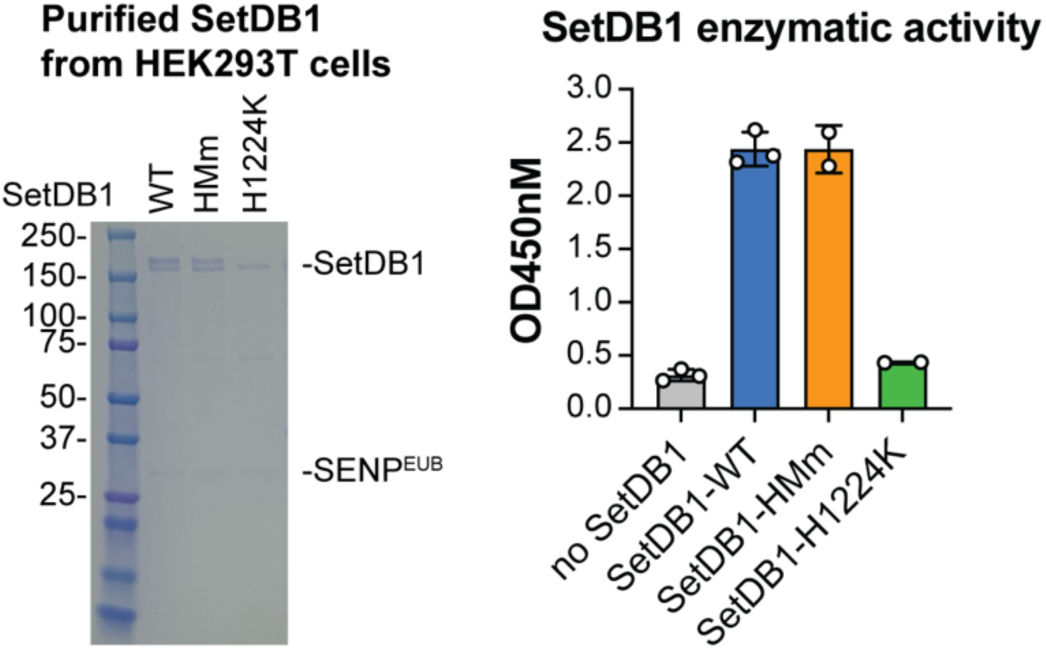
SetDB1-HMM mutation does not affect SetDB1 enzymatic activity *in vitro*. WT or mutant EGFP-SUMO^EU^-SetDB1 was expressed and purified from HEK293T cells. SetDB1 was eluted from the beads using SUMO protease SENP^EUB^ (left). Quantification of the methyltransferase activity of purified SetDB1 on histone H3 peptide was performed using a commercial kit. The amount of methylated H3K9 was quantified using an HRP-conjugated secondary antibody and color development system under OD450 (right). Dots represent independent biological replicates; bars indicate the mean ± SD.

**Fig. S9.**
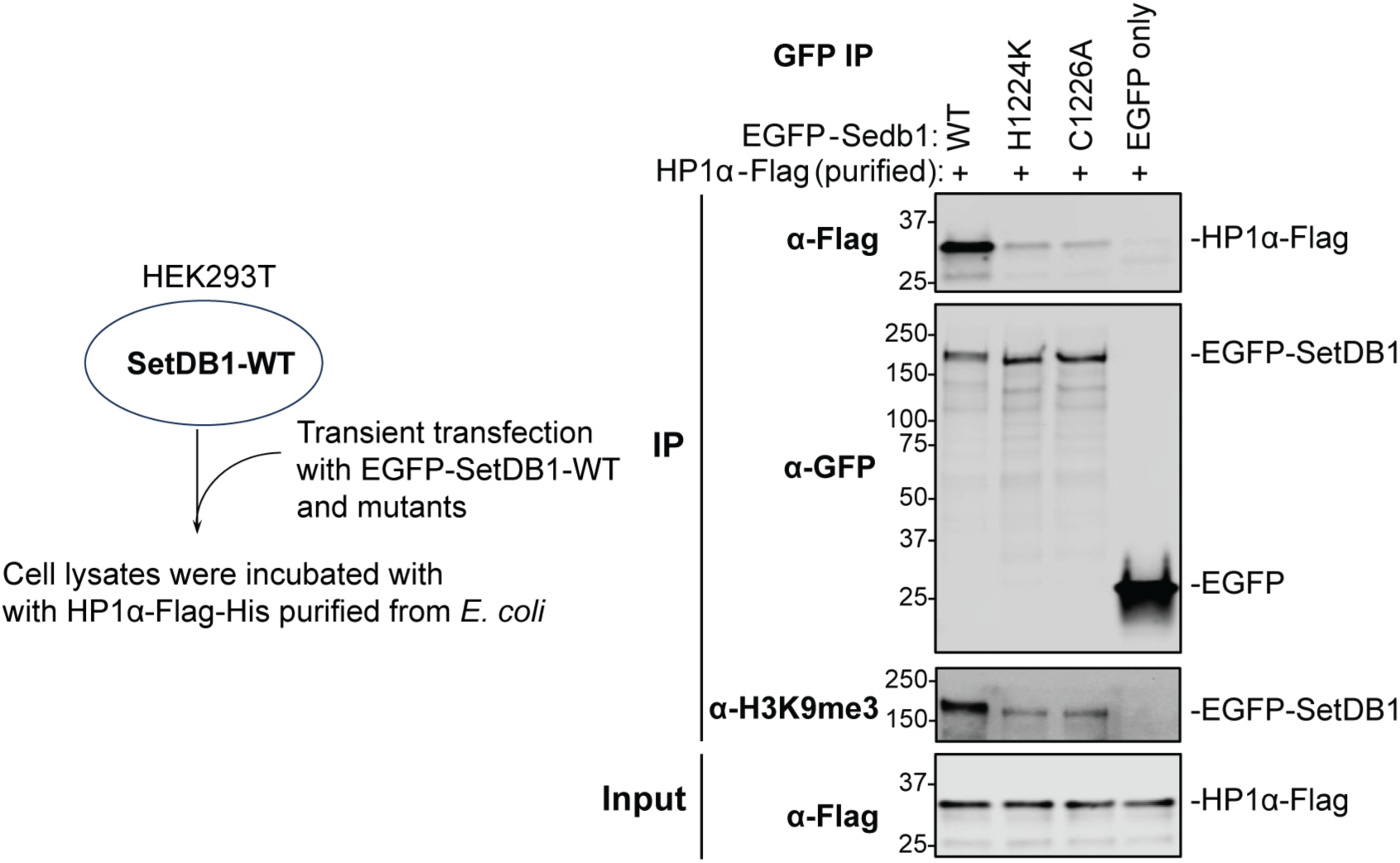
SetDB1 auto-methylates its histone mimic motifs *in cis*. EGFP-SetDB1-WT or methyltransferase-dead mutants (H1224K or C1226A) were transiently expressed in HEK293T cells that endogenously express SetDB1. Cell lysates were incubated with HP1α-Flag-His purified from *E. coli*, followed by co-immunoprecipitation using GFP nanotrap beads and western blot analysis. EGFP alone was used as a negative control. Both EGFP-SetDB1 methylation and its interaction with HP1α were strongly reduced in the mutants.

**Fig. S10.**
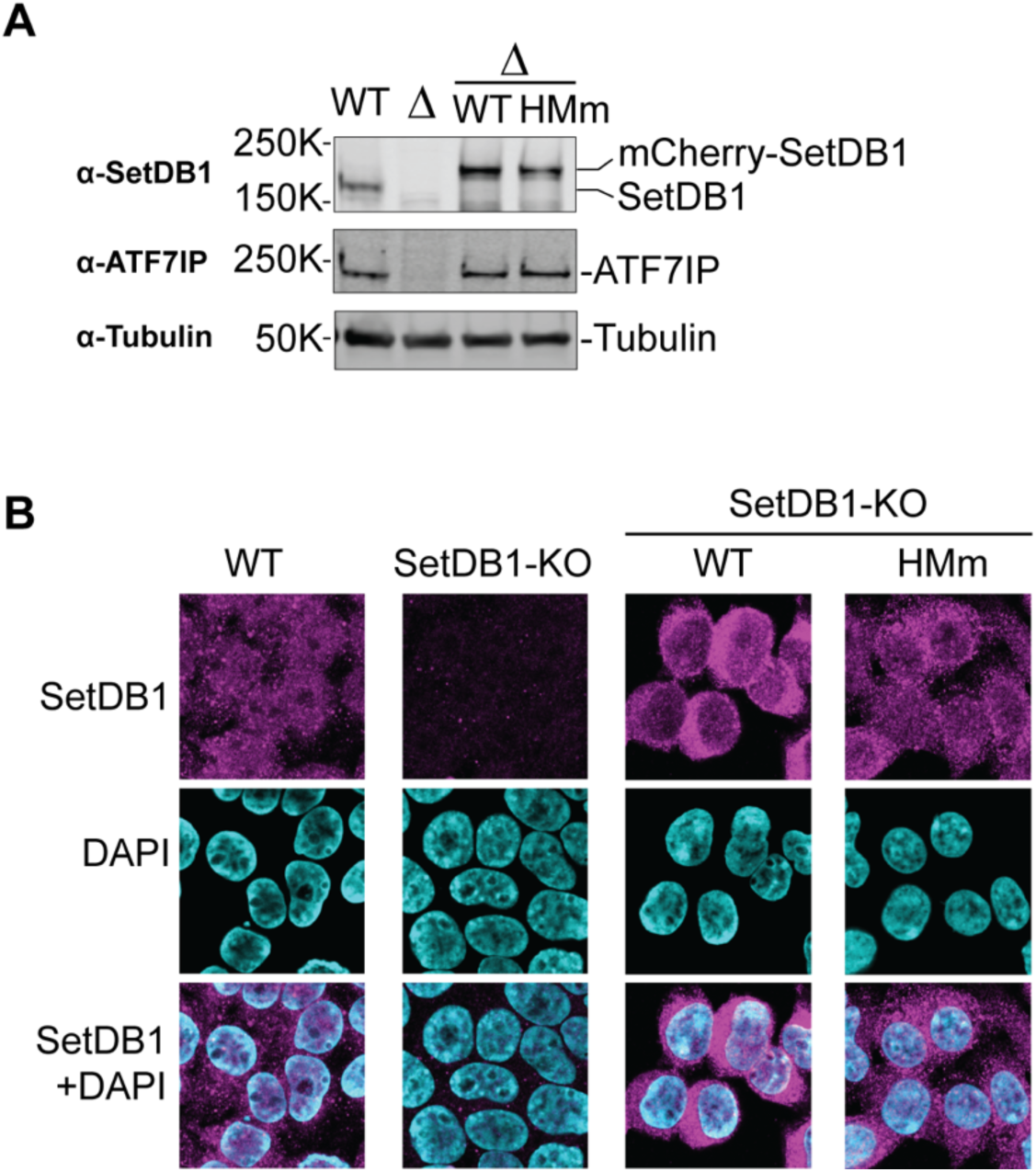
SetDB1-HMM mutation does not affect the nuclear localization of SetDB1. (A) SetDB1-HMm stabilizes the SetDB1 cofactor ATF7IP. WT or HMm mCherry-SetDB1 was stably expressed in SetDB1-KO HEK293 cells. Proteins were detected from total cell lysates by Western blot using antibodies against SetDB1, ATF7IP, and Tubulin. (B) SetDB1-HMm is present in the nucleus. WT or HMm mCherry-SetDB1 was stably expressed in SetDB1-KO HEK293 cells. Immunofluorescence was performed using an antibody against SetDB1. Nuclei were stained with DAPI.

**Fig. S11.**
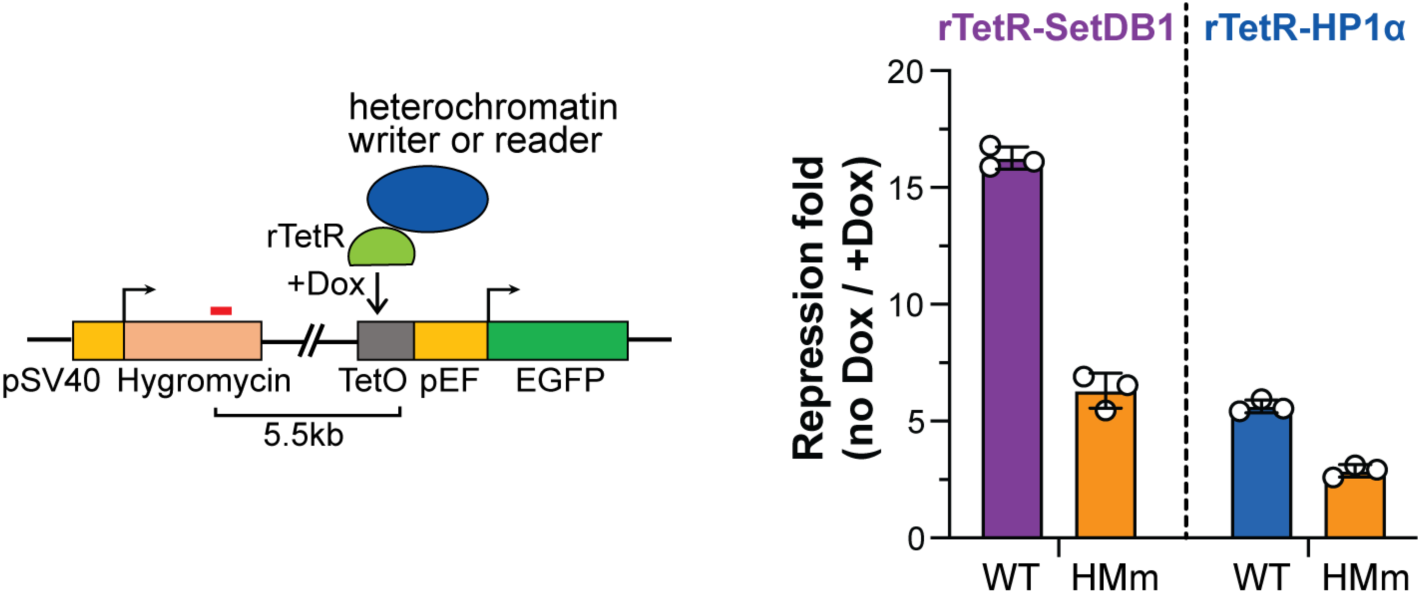
Loss of SetDB1 histone mimic methylation impairs silencing of the reporter-distal hygromycin resistance gene upon SetDB1 or HP1α tethering. WT or HMm SetDB1 was recruited to the reporter, or HP1α was recruited to the reporter in SetDB1-KO cells rescued with either WT or HMm SetDB1. The expression of the hygromycin resistance gene located 5.5 kb upstream of the tethering site was detected by RT-qPCR. GAPDH was used as an internal reference, and the repression fold was calculated as the fold difference in hygromycin expression in the absence or presence of Dox. The amplicon used for qPCR analysis is shown as a red bar.

**Fig. S12.**
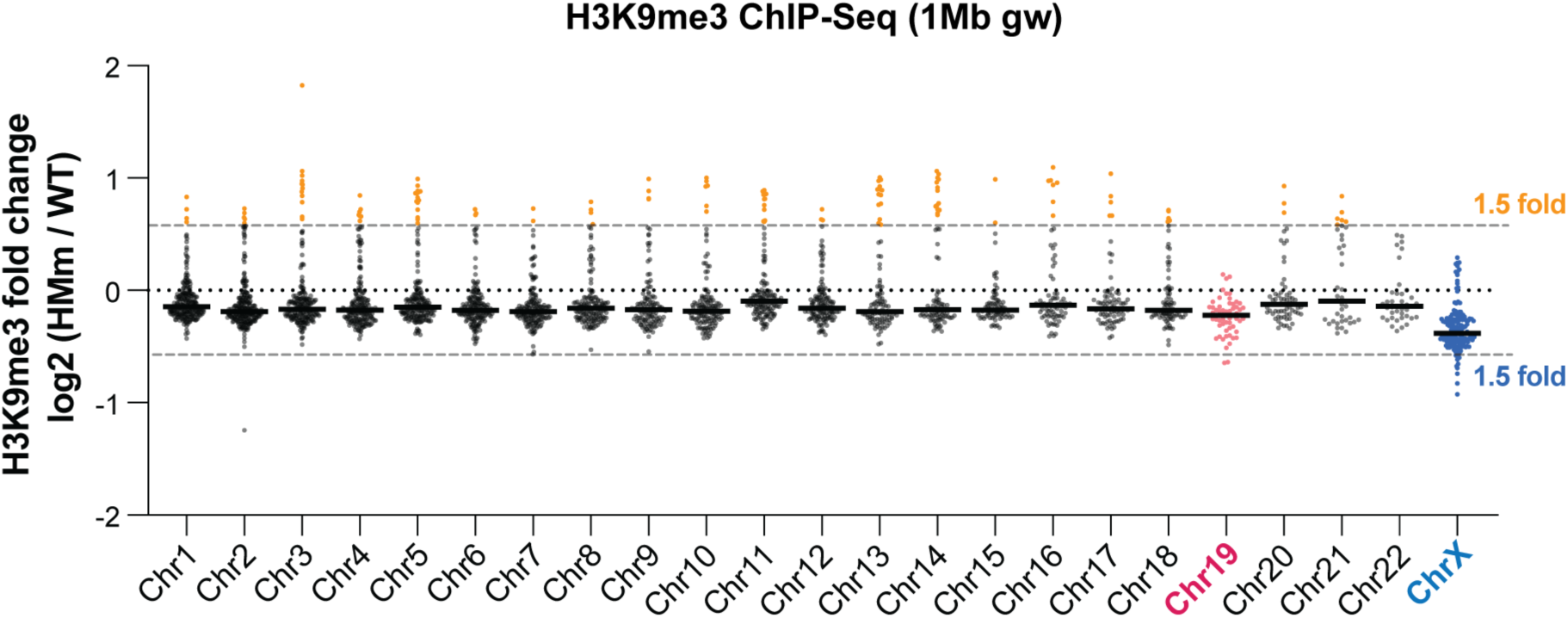
Regions with strong H3K9me3 signal gain in SetDB1-HMM mutant are scattered along all chromosomes except Chr19 and ChrX. H3K9me3 ChIP-seq was performed in SetDB1-KO cells in which SetDB1 loss was rescued by stably expressing either HMm or WT SetDB1. Dot plots show the fold change in H3K9me3 in SetDB1-HMm rescue strain compared to WT rescue strain in 1Mb genomic windows (gw) along the chromosomes. Gray dashed lines indicate a 1.5-fold (1.5x) change. Windows with >1.5x increase in H3K9me3 signal are shown in yellow.

**Fig. S13.**
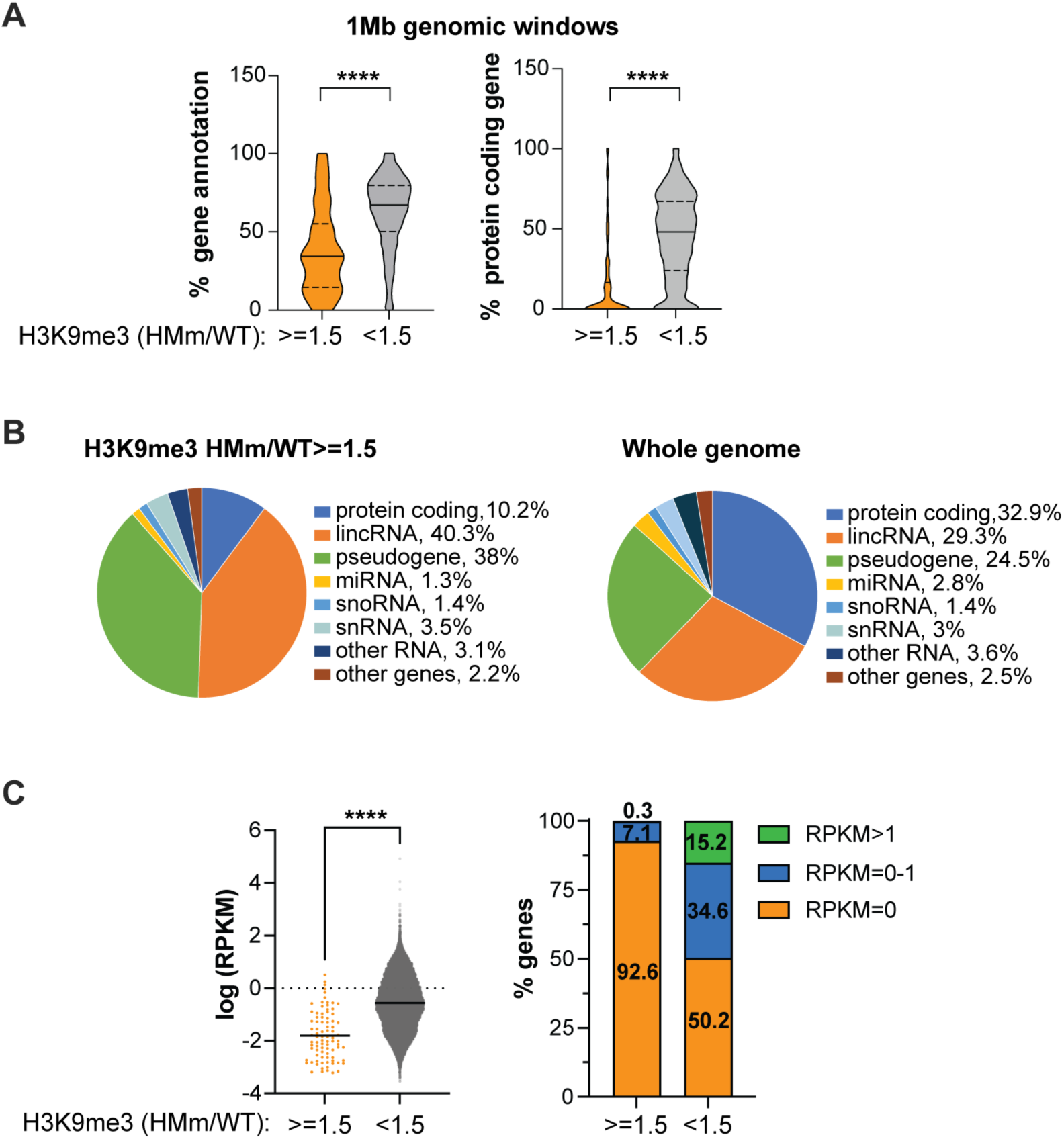
Regions with elevated H3K9me3 signals in SetDB1-HMM mutant are predominantly gene-poor and contain genes with low expression levels. (A) Violin plots show the proportion of gene (left) and protein-coding gene (right) annotations in the sequence of 1Mb genomic windows. Windows are categorized based on whether the H3K9me3 signal increases by ≥1.5x (yellow) or <1.5x (grey) in SetDB1-HMm rescue strain compared to WT rescue strain. The solid line indicates the median, and the dashed lines represent quantiles. p-value < 0.0001 is indicated by ****. (B) Gene type distribution in regions showing ≥1.5x elevated H3K9me3 signal in SetDB1-HMM mutant (left) and genome-wide (right). (C) Most genes in regions with ≥1.5x elevated H3K9me3 signal in SetDB1-HMM mutant show no or low expression. A dot plot (left) shows log-transformed RPKM values (from RNA-Seq of the WT rescue strain) for genes located in 1Mb windows with ≥1.5x or <1.5x increase in H3K9me3 signal in SetDB1-HMm rescue strain compared to WT rescue strain. The right panel shows the proportions of genes with different expression levels in both groups. Numbers indicate the percentage of genes within each group.

**Fig. S14.**
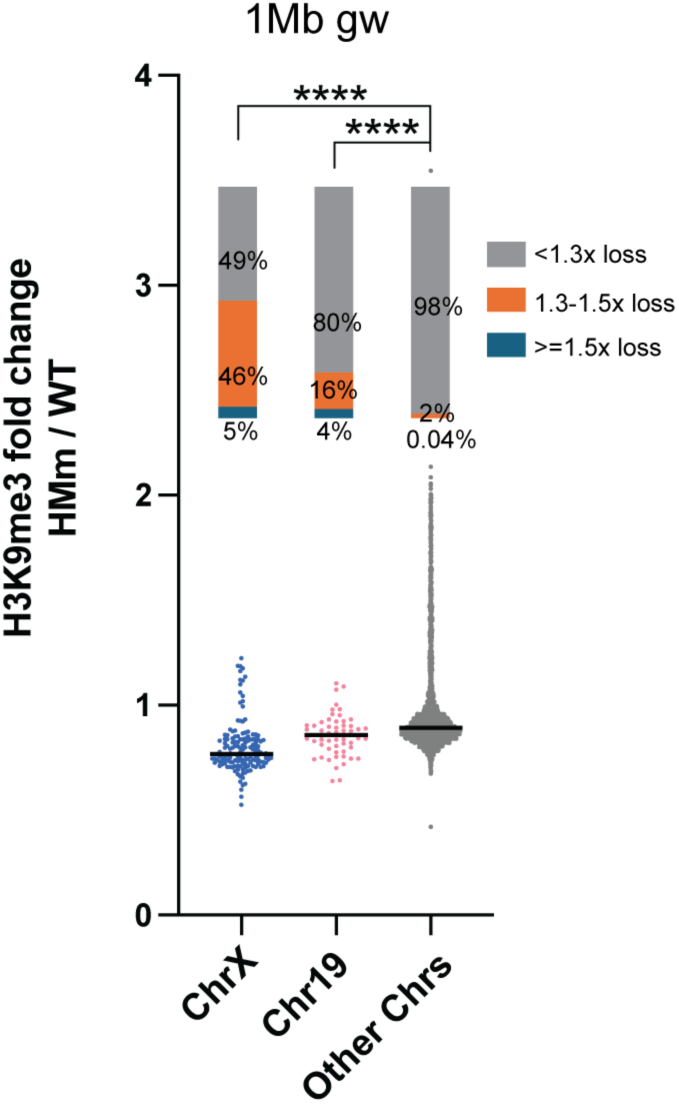
Regions with decreased H3K9me3 signal are mainly enriched on ChrX and Chr19. H3K9me3 signal decreased significantly more on ChrX and Chr19 compared to other chromosomes in SetDB1-HMm rescue strain compared to WT rescue strain. Dot plots (bottom) show fold changes of H3K9me3 (HMm/WT) in 1Mb genomic windows. The percentage of windows with ≥1.5x, 1.3-1.5x, or <1.3x loss of H3K9me3 in the mutant was calculated for each group. p-value < 0.0001 is indicated by ****.

**Fig. S15.**
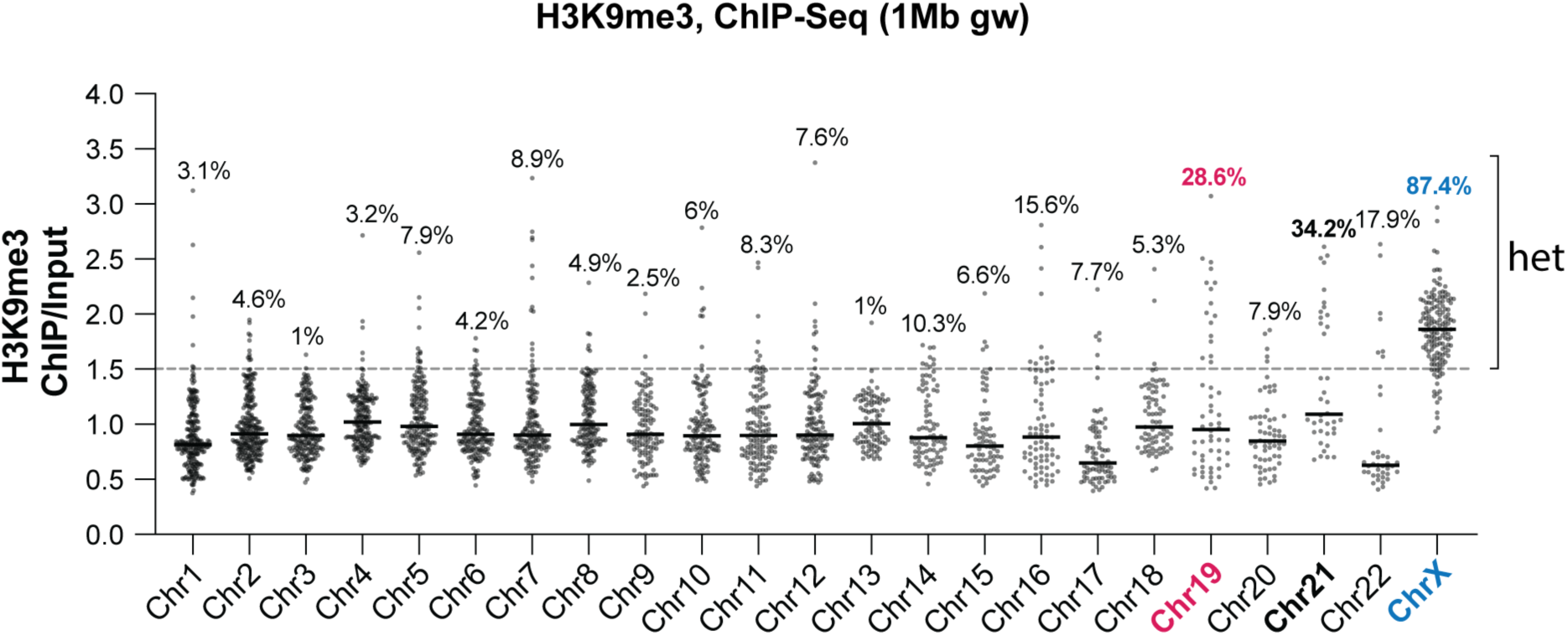
ChrX is heterochromatic in HEK293 cells. A dot plot shows H3K9me3 enrichment in 1Mb genomic windows across different chromosomes in SetDB1-WT rescue strain. Numbers indicate the percentage of heterochromatic windows (H3K9me3 ChIP/Input >1.5) for each chromosome.

**Fig. S16.**
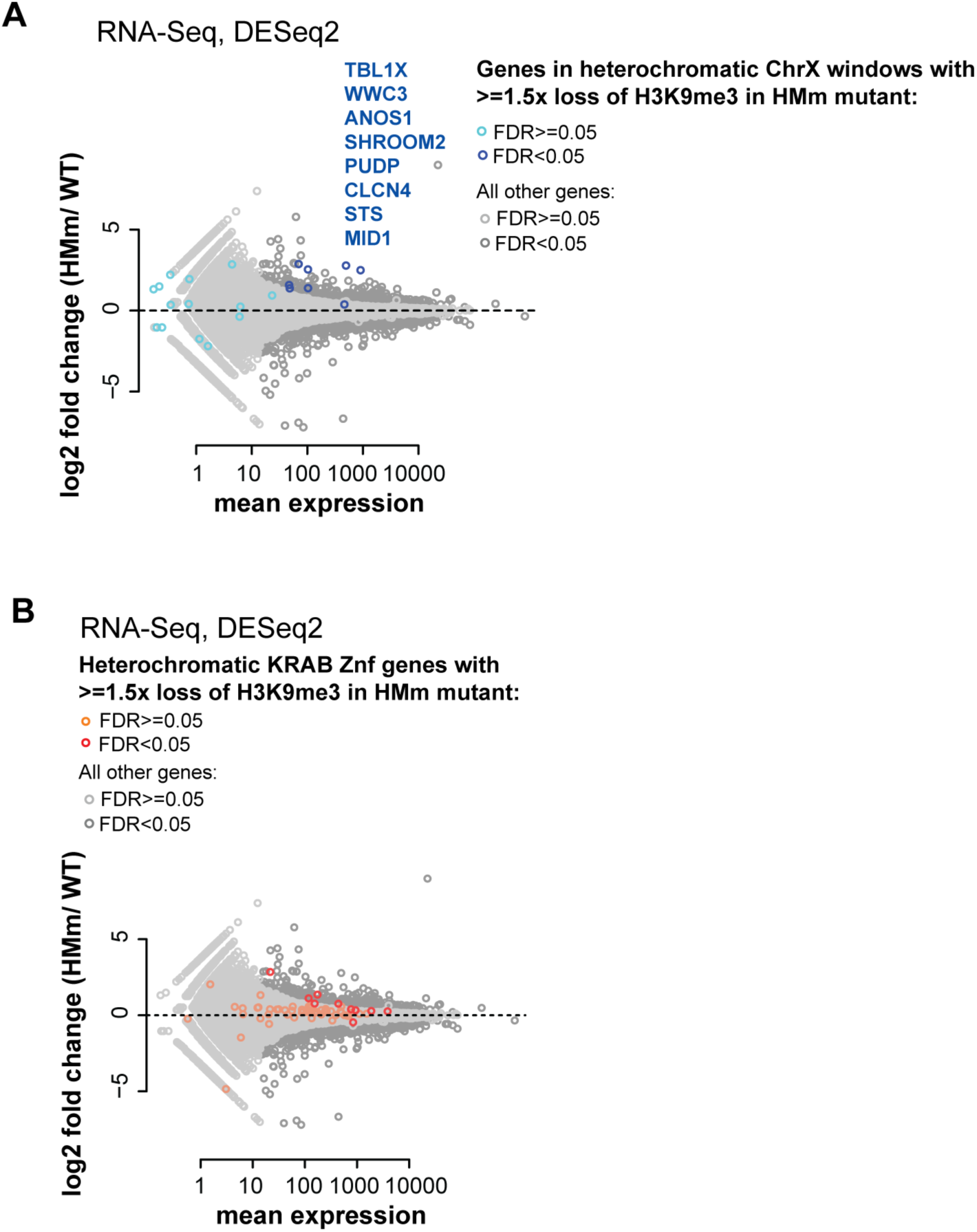
H3K9me3 loss on ChrX and over KRAB-Znf genes leads to mildly increased gene expression. (A) H3K9me3 loss correlates with increased gene expression on ChrX. RNA-seq data were analyzed by DESeq2 using 3 replicates for WT and 2 replicates for SetDB1-HMM mutant. ChrX genes located in 1Mb heterochromatic windows with ≥1.5x loss of H3K9me3 in the mutant are shown in blue; all other genes are shown in grey. ChrX genes with significantly increased expression in the mutant (FDR < 0.05) are listed. (B) Heterochromatic KRAB-Znf genes that show ≥1.5x loss of H3K9me3 exhibit mildly increased gene expression in SetDB1-HMm rescue strain. KRAB-Znf genes with ≥1.5x loss of H3K9me3 in the mutant are shown in orange (FDR ≥ 0.05) or red (FDR < 0.05), while all other genes are shown in grey.

**Fig. S17.**
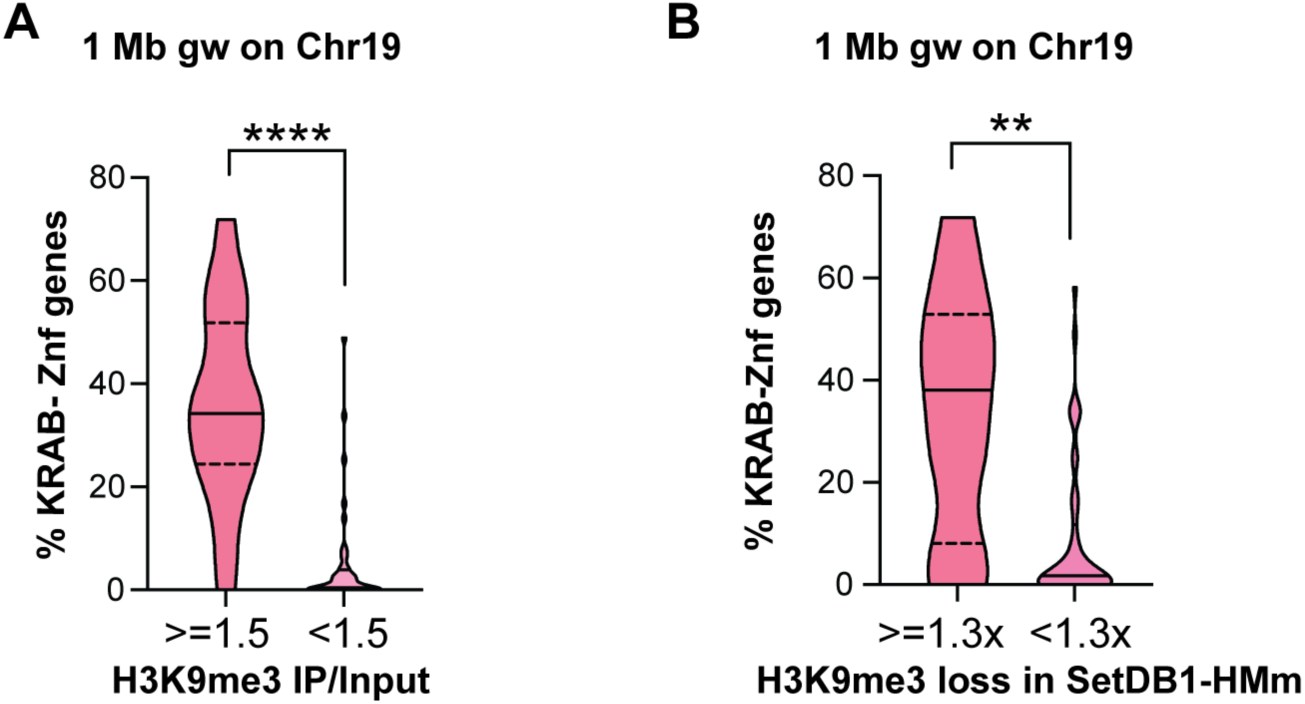
Regions with decreased H3K9me3 signals on Chr19 are enriched in KRAB-Znf genes. (A) KRAB-Znf genes on Chr19 reside in heterochromatin. Violin plots show the percentage of KRAB-Znf gene annotations within 1Mb windows on Chr19. Windows are categorized based on whether they are heterochromatic (H3K9me3 ChIP/Input ≥1.5 in WT rescue strain) or euchromatic (H3K9me3 ChIP/Input <1.5 in WT rescue strain). p-value < 0.0001 is indicated by ****. (B) Regions on Chr19 that lose H3K9me3 signal in SetDB1-HMm strain are enriched in KRAB-Znf genes. Violin plots show the percentage of KRAB-Znf gene annotations within 1Mb windows on Chr19. Windows are categorized based on whether H3K9me3 signal loss in SetDB1-HMm rescue strain was ≥1.3x or <1.3x. p-value < 0.01 is indicated by **.

